# Cooperation and spatial self-organization determine ecosystem function for polysaccharide-degrading bacteria

**DOI:** 10.1101/640961

**Authors:** Ali Ebrahimi, Julia Schwartzman, Otto X. Cordero

**Affiliations:** Ralph M. Parsons Laboratory for Environmental Science and Engineering, Department of Civil and Environmental Engineering, Massachusetts Institute of Technology, Cambridge, MA 02139 USA

**Author notes:** equal contribution (Ali Ebrahimi, Julia Schwartzman).

**Keywords:** marine microbes, cooperation, particulate organic matter, public goods, spatial organization

## Abstract

The recycling of particulate organic matter (POM) by microbes is a key part of the global carbon cycle, one which is mediated by the extracellular hydrolysis of polysaccharides and the production of public goods that can trigger social behaviors in bacteria. Despite the potential importance of these microbial interactions, their role in regulating of ecosystem function remains unclear. In this study, we developed a computational and experimental model system to address this challenge and studied how POM depolymerization rate and its uptake efficiency –two main ecosystem function parameters– depended on social interactions and spatial self-organization on particle surfaces. We found an emergent trade-off between rate and efficiency resulting from the competition between oligosaccharide diffusion and cellular uptake, with low rate and high efficiency being achieved through cell-to-cell cooperation between degraders. Bacteria cooperated by aggregating in cell-clusters of ~10-20μm, where cells were able to share public goods. This phenomenon, which was independent of any explicit group-level regulation, led to the emergence of critical cell concentrations below which degradation did not occur, despite all resources being available in excess. By contrast, when particles were labile and turnover rates were high, aggregation promoted competition and decreased the efficiency of carbon utilization. Our study shows how social interactions and cell aggregation determine the rate and efficiency of particulate carbon turnover in environmentally relevant scenarios.

**Significance Statement:** Microorganisms can cooperate by secreting public goods that benefit local neighbors, however, the impact of cooperation on ecosystem functions remains poorly constrained. We here pair computation and experiment to show that bacterial cooperation mediates the degradation of polysaccharide particles recalcitrant to hydrolysis in aquatic environments. On particle surfaces, cooperation emerges through the self-organization of cells into ~10-20μm clusters that promote cooperative uptake of hydrolysis products. The transition between cooperation and competition in aggregates is mitigated by individual cell behaviors such as motility and chemotaxis, that promote reorganization on the particle surface. When cooperation is required, the degradation of recalcitrant biopolymers can only take place when degraders exceed a critical cell concentration, underscoring the importance of microbial interactions for ecosystem function.

## Introduction

The microbial breakdown of complex polysaccharides is a key ecosystem process that enables the recycling of carbon from plant and animal detritus into global biogeochemical cycles and is a relevant process in all heterotrophic microbial ecosystems, from animal guts (1–3) to soils (4, 5) and oceans (6–8). A key feature of these polysaccharides is their insoluble nature: a large fraction is found in particles at the scale of 100 μm, that require both surface colonization and extracellular hydrolysis to be degraded (9, 10). On particle surfaces cells can attach and grow in close proximity, increasing the opportunity for microbial interactions to impact the ecosystem process. One particularly relevant type of interaction in this context is cell-cell cooperation mediated by the sharing of public goods such extracellular enzymes and hydrolysis products (11). However, the extent to which these interactions take place and impact bacterial growth in the environment remains unclear. Previous work on cooperative interactions has largely focused on the opportunity that public goods open for exploitative populations to invade (so-called cheaters), and less on the environmental and physiological conditions that enable cooperation to take place or the potential role of cooperative behavior on ecosystem processes. From the perspective of ecosystem modeling, efforts to incorporate the role of microbes in organic matter degradation equate microbial activity with enzymatic activity without considering the role of population-level phenomena such as cooperation. In this paper we reveal how microbial social interactions can impact relevant ecosystem parameters.

The extent to which social interactions mediated by public goods play a relevant role in ecosystem function is highly dependent on how public goods diffuse (12, 13). In a three-dimensional aqueous environment like the ocean, if cells are too far apart only a minuscule fraction of the public goods are recovered by neighbors, while the rest is lost to the environment. In contrast, if cells are sufficiently proximal to each other and the resource is limiting, growth kinetics can be cooperative, meaning that the per-capita growth rate is positively dependent on the density of degrader cells (14). This logic suggests that cooperation should be accompanied by the emergence of spatial patterns, such as cell patches. If the cooperative effects in these patches are strong, critical population density thresholds might emerge below which degradation cannot support population growth (14, 15). Less recognized is the contribution of individual cell behaviors, such as surface attachment, chemotaxis, and biofilm formation, on the ability of cells to find those critical densities by aggregating into cell patches. Therefore, in order to begin to understand the role of social interactions in natural systems, we need to take into account the physical constraints of the micro-environment and how populations interact with these constraints through their behavior.

To quantify the impact of bacterial social interactions and spatial behavior on ecosystem function, we focus on two main parameters: the speed at which polymers are hydrolyzed and converted to soluble oligosaccharides, that is, the turnover rate (16, 17), and the POM uptake efficiency, which is the fraction of the dissolved oligosaccharide that can be taken up by cells and converted into biomass. To study the role of social interactions and spatial behavior on ecosystem function, we developed a computational and experimental model of the colonization of insoluble particulate polysaccharides by marine heterotrophic bacteria. The individual-based model (18, 19) simulates the functional traits of individual cells: chemotactic movement, particle attachment and detachment, the secretion of enzymes, oligosaccharide uptake and growth. The experimental system validates computational predictions in a chitin-degrading bacterial strain isolated from the coastal ocean, and clarifies the role of physiological parameters on social interactions (17). We leveraged the computational model to study the relationship between degradation rate and POM uptake efficiency and how emergent bacterial behaviors influence their ability to degrade recalcitrant particles, and we tested some of our predictions using our experimental model of chitin colonization. Our work demonstrates that cell-cell cooperation is critical for the degradation of complex biomaterials, implying that the degradation of recalcitrant polysaccharides can be bacteria-limited. Moreover, cell-density thresholds that determine the onset of cooperative growth depend strongly on individual cell behavior, in particular those behaviors that regulate the residence time of bacteria on particles.

## Results

We modeled the dynamics of cell colonization, enzyme secretion and growth (Figure 1A) using an individual based model to describe cells, coupled to a reaction-diffusion framework to describe enzymes and oligosaccharides. In the model, bacterial cells that attached to the surface of a polysaccharide particle broadcast enzymes that reacted with the surface of the particle, releasing oligosaccharides to which non-attached cells could chemotax. Cellular uptake of oligosaccharides followed Monod kinetics (20) and cells were allowed to divide after a certain quota of oligosaccharide is consumed (19) (see Methods and Supplementary Information for a detailed description and Table S1 for the parameters). This individual-based approach allowed us to modulate traits such as chemotaxis or particle-attachment rate, and measure their impact on the carbon uptake rate on a cell by cell basis.

**Figure 1.**
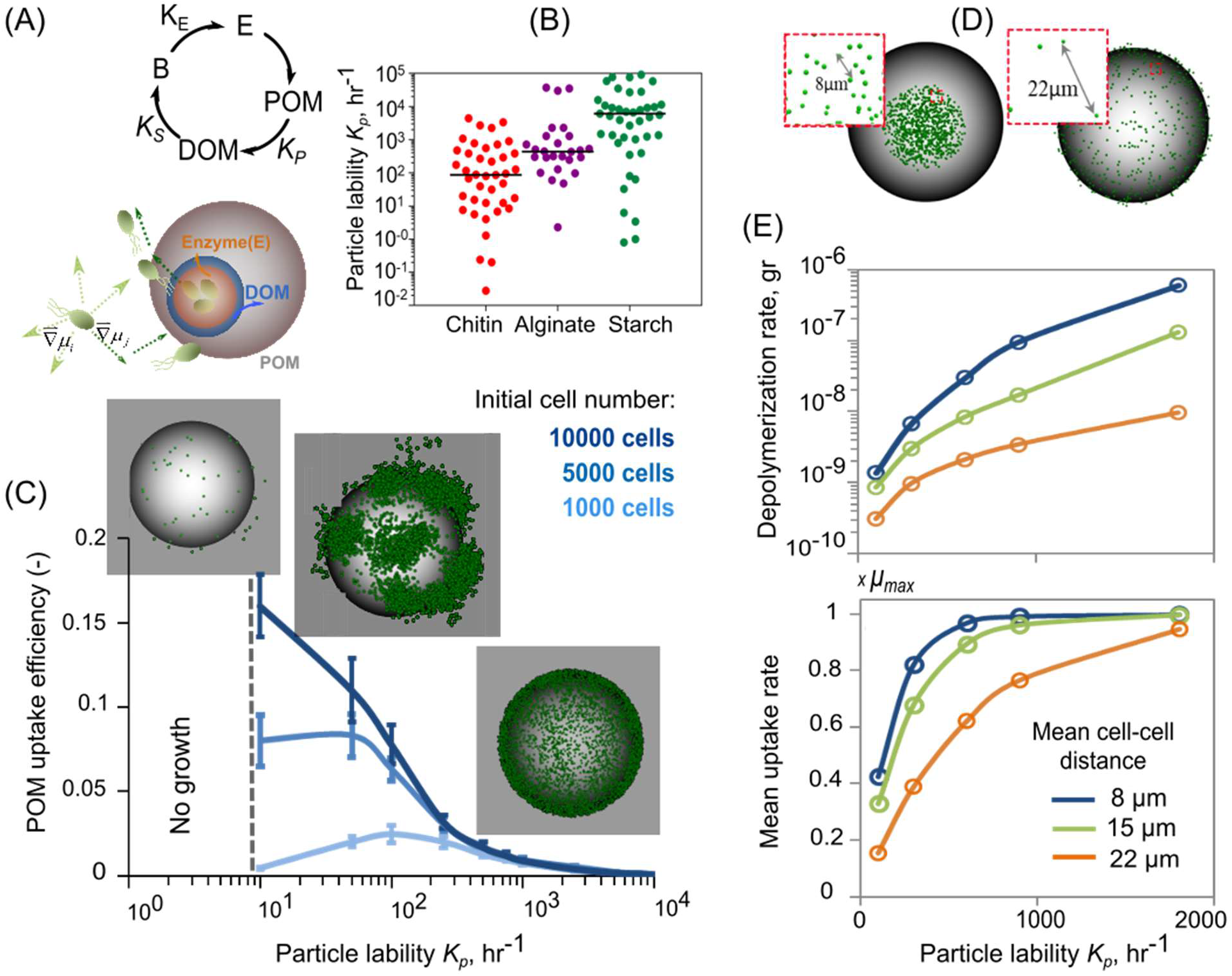
POM uptake efficiency is regulated by an emergent rate-yield trade-off. A) Conceptual representation of enzyme secretion by bacterial cells, breakdown of polysaccharide substrates to oligosaccharides by enzymes and uptake of oligosaccharides by bacterial cells. Schematic representation of trait based modeling of microbial dispersal and colonization on the particle is shown (below). B) The distribution range of particle lability (K_p_) from natural polymeric carbohydrates (Chitin, Alginate, Starch). The data are for various bacterial species with their corresponding abiotic conditions (species name, substrates and environmental conditions are represented in supplementary Table S2) (data are from Brenda database: www.brenda-enzymes.org). The solid line indicates the mean value of the particle lability. C) POM uptake efficiency as a function of particle lability and initial population size. Dashed line indicates the no growth zone. Microbial population assembly on the particle for three levels of particle lability (recalcitrant I: K_p_:1hr^−1^, semi-labile II: K_p_:100hr^−1^ and labile III: K_p_:1000hr^−1^) are shown. Green dots show individual cells on the particle. Simulations are performed for a range of initial cell densities and 1% detachment is allowed. Half saturation, K_s_ is assumed 0.1mg/L. D) The cell spatial distribution on particles for scenarios with 8 and 22μm mean cell distances are shown. E) Depolymerization and mean uptake rates for a range of mean cell-cell distance are represented as a function of particle lability. The simulations are initialized with placing individual cells uniformly on the particle to meet their corresponding mean cell-cell distance for neighboring cells. No detachment is allowed in simulations. Initial cell number was 1000 cells.

A crucial parameter of our model was the “particle lability”, *K*_*p*_, which defined how many grams of oligosaccharide were released per gram of enzyme acting on the polysaccharide surface per unit of time. *K*_*p*_ was a compound parameter that resulted from the product of the catalytic activity of the enzyme, *k*_*cat*_ and the recalcitrance of the substrate. This parameter played a central role because it determined the maximum degradation rate and controlled the nutrient supply rate of bacteria. A survey of hydrolysis rate values reported in the literature revealed that the particle lability, *K*_*p*_ can exhibit significant variation across natural environments and microbial enzymes. *K*_*p*_ varied by more than 6 orders of magnitude within glycosyl hydrolase families- a trend that held true among different substrate types such as chitin, alginate and starch (Figure 1B). This led us to ask how variation in particle lability, *K*_*p*_ affected population growth dynamics and the rate – efficiency relation of POM depolymerization.

Our results revealed that there is an emergent negative relationship between the rates of depolymerization and growth, and the POM uptake efficiency of the particle-associated bacterial population (Figure 1C) (21, 22). This emergent rate – efficiency trade-off was a consequence of the diffusion of oligosaccharide in a three dimensional environment where soluble products that were not taken up by cells in the vicinity of the particle were lost. At high values of *K*_*p*_, oligosaccharides are produced in excess of *K*_*s*_, the half-saturation constant of the Monod growth function, and therefore cells approached their maximum growth rate. However, high *K*_*p*_ also led to ~99% loss of oligosaccharide to diffusion (~1% recovery), which reduced the theoretical biomass yield of the population and the POM uptake efficiency. For comparison, if the system was closed, as in a laboratory reactor, POM uptake efficiency could theoretically reach 100% because dissolved oligosaccharides would accumulate (Figure S1). However, natural environments are rarely, if ever, closed and diffusive losses are likely to limit POM uptake efficiency in nature, given low particle densities, (Table S3). Moreover, adding the movement of fluids around particles by convective flow to the model (23–25), further increased the loss rate of oligosaccharide to the bulk environment, and reduced POM uptake efficiency from 10% to 2% (Figure S2–S5). Although the exact value of POM uptake efficiency could depend on substrate affinity (1/*K*_*s*_) and on cell numbers (the more cells that can capture oligosaccharides the higher uptake efficiency), the trade-off between rate and efficiency held for different physiological parameters (Figure S6). Taken together, these results suggest that in natural environments most public goods are lost and that the competition between diffusion and uptake should lead to a tradeoff between POM uptake efficiency and its turnover rate. Quantifying the uptake efficiency for natural marine particles revealed that the maximum efficiency barely exceeded 7% at the optimum particle lability (*K*_*p*_ ~100hr^−1^) for the highest of particle-associated cell density observed (~2.32×10^7^) (Table S3).

Surprisingly, we found that the high POM uptake efficiency observed at low *K*_*p*_ (recalcitrant particles/low enzymatic activity per cell) was mediated by the aggregation of cells into micro-scale patches on the particle surface, a phenomenon that was not hardcoded in the model but emerged from the interplay between diffusion, cell behavior and growth (Figure 1C and Figure S7). Within these patches, cells grew cooperatively by sharing oligosaccharides that would otherwise be lost to diffusion, which increased the per capita growth rate and POM uptake efficiency up to a density of 0.3 cells/μm^2^ (Figure S8). To characterize the spatial density dependence, we performed simulations to quantify particle depolymerization and mean growth rates as a function of the inter-cell distance (Figure 1D). Our analysis showed that dense spacing (a nearest neighbor distance of 8 μm) promoted cooperation by sharing of oligosaccharides, but only when particles were recalcitrant and the oligosaccharide production rate was slow (*K*_*p*_<100hr^−1^) (Figure 1E). More precisely, when the amount of oligosaccharide available to cells fell near *K*_*s*_, the half-saturation of the Monod growth curve, an increase in the local concentration of oligosaccharide due to cell-cell aggregation increased the per capita growth rate. In contrast, at high *K*_*p*_ (~2000 hr^−1^), oligosaccharides quickly accumulated and the uptake rate was decoupled from the spatial organization of the cells on the particle, since there were enough resources for cells to grow at their maximal rate ([C]≫*K*_*s*_) (Figure 1E). Under these conditions, there is no benefit to aggregation and even cells spaced 22 μm apart reached their maximum oligosaccharide uptake rate (Figure 1E).

In our model, cell detachment and reattachment from the particle surface was a critical behavior that enabled the formation of patches and the degradation of recalcitrant particles. On recalcitrant particles (Kp=10-100 hr-1) 1% detachment significantly increased the particle degradation rate and its uptake efficiency (Figure 2A), and also increased the mean carbon uptake rate by a factor of 5 (Figure S9A), compared to a non-detaching population. This allowed populations to survive on recalcitrant particles that might otherwise not sustain growth and drive the population to extinction (Figure S9B). Without chemotaxis, random motility alone still allowed detaching populations to grow on more recalcitrant particles than non-detaching populations, but at ~1/6 the POM uptake efficiency (Figure 2A) and ~1/10 the rate of biomass accumulation (Figure S9B). This was due to the fact that with chemotactic motility most cells had access to the same of hydrolysis products emanating from cell patches, with the distribution of carbon uptake rates for individual cells displaying a tight peak near the maximum uptake rate (μ~0.8μmax) (Figure S9C). Our model thus suggests that detachment and chemotaxis enhance POM uptake efficiency under nutrient-limited conditions ([C]~ Ks) by enabling the formation of patches where cells cooperate by sharing public goods. This also implies that the onset of cooperation is dependent on the individual strain physiology. Organisms that had either a high affinity for oligosaccharides (low Ks) or a high hydrolytic activity saturated their growth at low oligosaccharide concentrations (Figure 2B), circumventing the need to cooperate Figure (2C-D). In contrast, organisms with a low uptake affinity for the public good, or organisms with a low per-cell rate of hydrolysis, such as those that tether enzymes to their membrane, had a higher need to cooperate with other cells (Figure 2D). Therefore, although there was a general trend to increase cooperation as particles became harder to degrade (Figure 2D), traits such as motility, surface detachment rates, substrate affinity or enzyme localization determined determine the exact onset of cooperation for each population.

**Figure 2.**
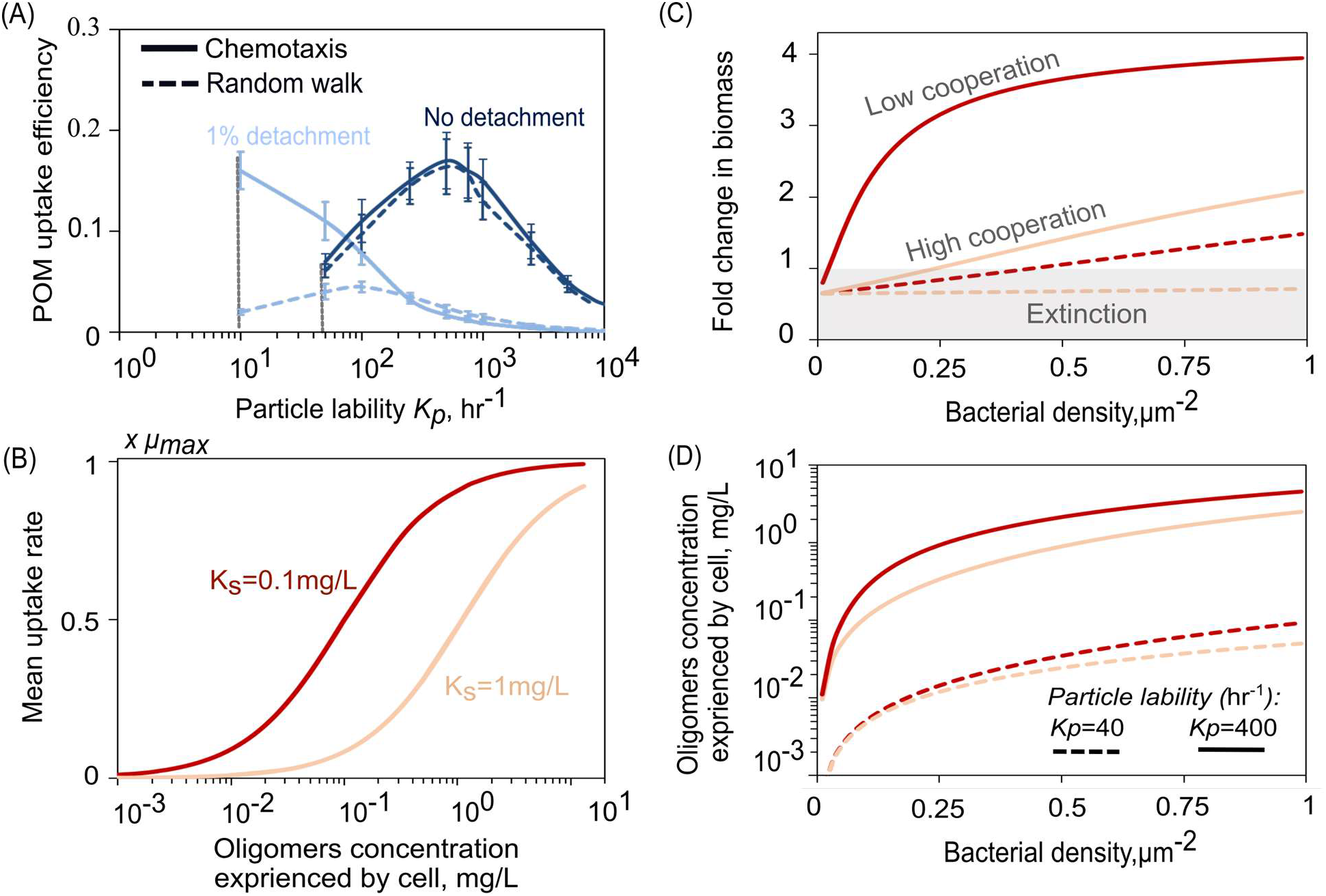
Bacterial cooperation enhances carbon uptake rate. A) Effects of individual cells attachment/detachment frequency to/from particles on POM uptake efficiency. Solid line shows the simulation results with chemotactic behavior compared to simulations with only random walk shown with dashed line. The computational results are shown for the initial cell number of 10^7^cells/ml and the particle size was set to 200 μm. The results are shown for simulations after 10 hours. No bacterial growth is expected for K_p_ values below the grey dashed line. B) Monod growth kinetics represents the mean cellular uptake rate of oligosaccharides as a function of oligosaccharide concentration. C) The fold of change in biomass as a function of initial bacterial density is shown for simulations of bacterial colonization on a single particle after 20hours. D) The effects of bacterial cell density on produced oligosaccharide concentration, experienced by individual cells. The simulations are performed for a particle with constant radius of 200 μm and initial cell density of 0.3 cell/μm^−2^. Labile (K_p_=400 hr^−1^) vs. recalcitrant (K_p_=40hr^−1^) particles are considered with two relatively low (K_s_=1mg/L) and high (K_s_=0.1mg/L) affinity to substrates uptake by bacterial cells.

To experimentally validate our prediction that cell-cell cooperation drives the degradation of hard-to-degrade polysaccharides, we turned our attention to *Psychromonas* sp., *psych6C06*, a marine isolate that had previously been enriched from coastal seawater on model chitin particles(10). The strain readily degrades chitin hydrogel in ~30 hours (17) and encodes at least eight predicted chitinases, or glycosyl hydrolase family 18 and 19 homologs, but no other families of glycosyl hydrolases, leading us to conclude that the strain is representative of a chitin specialist. We reasoned that if cooperative growth kinetics played a role in this system, we would observe a strong dependency between the initial number of cells that can colonize the particle and the growth of the population. In particular, we would expect a critical cell density below which the population is unable to form the patches required to degrade particles, revealing that the degradation process is bacteria-limited.

In agreement with this prediction, *psych6C06* displayed a strong density dependence when growing on hydrogel chitin beads, in the form of a critical cell density below which degradation never occurred (Figure 3A). Interestingly, we observed that colonization involved the formation of cell patches, in agreement with the model results (Figure 3B). At concentrations just below the threshold critical cell density, we saw that cells that initially colonized the particle were not able to persist. Populations that persisted did so by forming cell patches (Figure 3C). We artificially increased *K*_*p*_ by adding exogenous chitinase to supply 776 μg/h GlcNAc. Consistent with individual-based model results (Figure 2D), the addition of the exogenous enzyme activity lowered the cell density-dependent threshold for colonization of the chitin hydrogel beads (Figure 3D). In addition, the broadcast chitinases led to a more uniform distribution of *psych6C06* cells on the chitin hydrogel bead, a state that was morphologically distinct from the patchy colonization achieved at 24 h *psych6C06* without enzyme (Figure 3B, Figure S10C) and similar to the simulation results at high *K*_*p*_ (Figure 2D).

**Figure 3.**
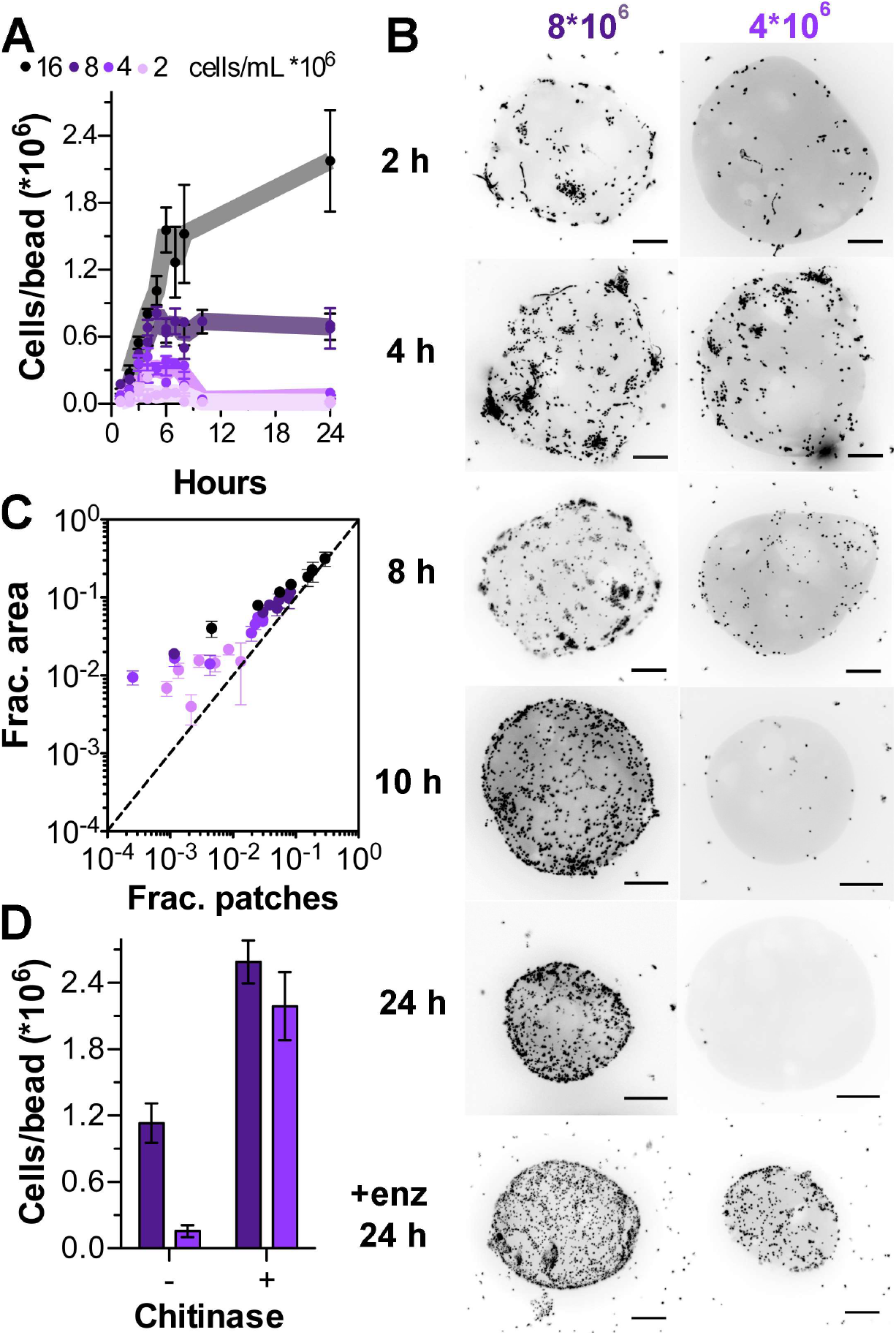
Chitinase limitation drives psych6C06 to form growth-promoting surface-associated clusters. A) An Allee effect emerges for populations of psych6C06 over 24 h. Data points are combined from three experimental replicates. Error bars are SEM from at least 6 individual measurements of colonization density on chitin beads. Lines represent the mean trajectory for each cell density. B) Representative images showing colonization density of initial populations of cells below (4*10^6^ cells/mL), and above (8*10^6^ cells/mL) the colonization threshold at timepoints during colonization. Scale bars are 20 μm. C) The fraction of cell area (Frac. area) on chitin hydrogel beads that exists in patches >2000 cells (Frac. patches). Dashed line indicates the limit where all cell area exists in patches (x=y). Raw data analyzed are the same as 3A and 3B. D) The addition of exogenous chitinase enables smaller populations of cells to colonize chitin hydrogel particles. Dark purple bars, initial density 8*10^6^ cells/mL; light purple bars, initial density 4*10^6^ cells/mL. Colonization density was assessed after 24 h. Bars are SEM from at least 5 measurements. The amount of chitinase added to the medium supplies 776 μg/h GlcNAc.

To obtain a more mechanistic understanding of the critical threshold phenomenon we took a bottom-up approach by predicting the threshold cell density based on measurements of the relevant physiological and behavioral parameters of *psych6C06*. We calculated particle attachment and detachment rates by quantifying cell density on particles (Figure 4A-B). Our measurements revealed rapid attachment and detachment rates (attachment rate 0.03 h^−1^, detachment rate 0.26 h^−1^) that were proportional to the density of cells off and on the particle, respectively suggesting that the population undergoes frequent rearrangement on the surface of the particle. These observations echo the rearrangement observed in the individual based simulations (Figure 2A). We measured *K*_*p*_ for *psych6C06*, using the fluorescent substrate 4-Methylumbelliferyl (MUF)-*N*-acetyl-β-D-glucosaminide, which detects the release of GlcNAc from chitin. We noted that very little chitinase activity was detected in the culture supernatant of *psych6C06*, but robust activity was associated with the cells themselves (Figure 4C, Figure S10A) indicating that enzymes were membrane bound. Using MUF-conjugated substrates with different cleavage specificities, we determined that most of *psych6C06* chitinase activity was derived from exochitinase, which releases GlcNAc as a product (Figure S10B). Thus, we assessed the GlcNAc to biomass conversion factor (the biomass yield) of this strain by direct measurement of sugar consumption and cell density in exponentially growing cultures (Figure 4D). We measured of the growth rate of *psych6C06* on a range of GlcNAc concentrations, and used these substrate-limited growth measurements to derive *μmax* and *K*_*s*_ from a fit of the Monod growth equation(20) (Figure 4E). We used the diffusive loss of oligosaccharides predicted by our 3D simulations to estimate GlcNAc loss, and assumed that all cells on the surface experienced the same concentration (no local gradients). Using these measurements, we parameterized a simple version of the individual based model written to describe population-level growth dynamics on a surface where cells can attach, detach, and grow as a function of the hydrolyzed product concentration (see Methods for a full description of the analytical bottom-up model). Using this simplified model, with no free parameters, we studied how the initial cell density determines the population colonization rate and the growth of bacteria on the particle surface. We found a remarkable quantitative agreement between model and experiments, with critical thresholds predicted between initial densities of 5*10^6^ and 10^7^ cells/mL (Figure 4F). Our analysis thus shows that individual cell level physiology and behavior together with diffusion regulate the onset of social degradation of POM.

**Figure 4.**
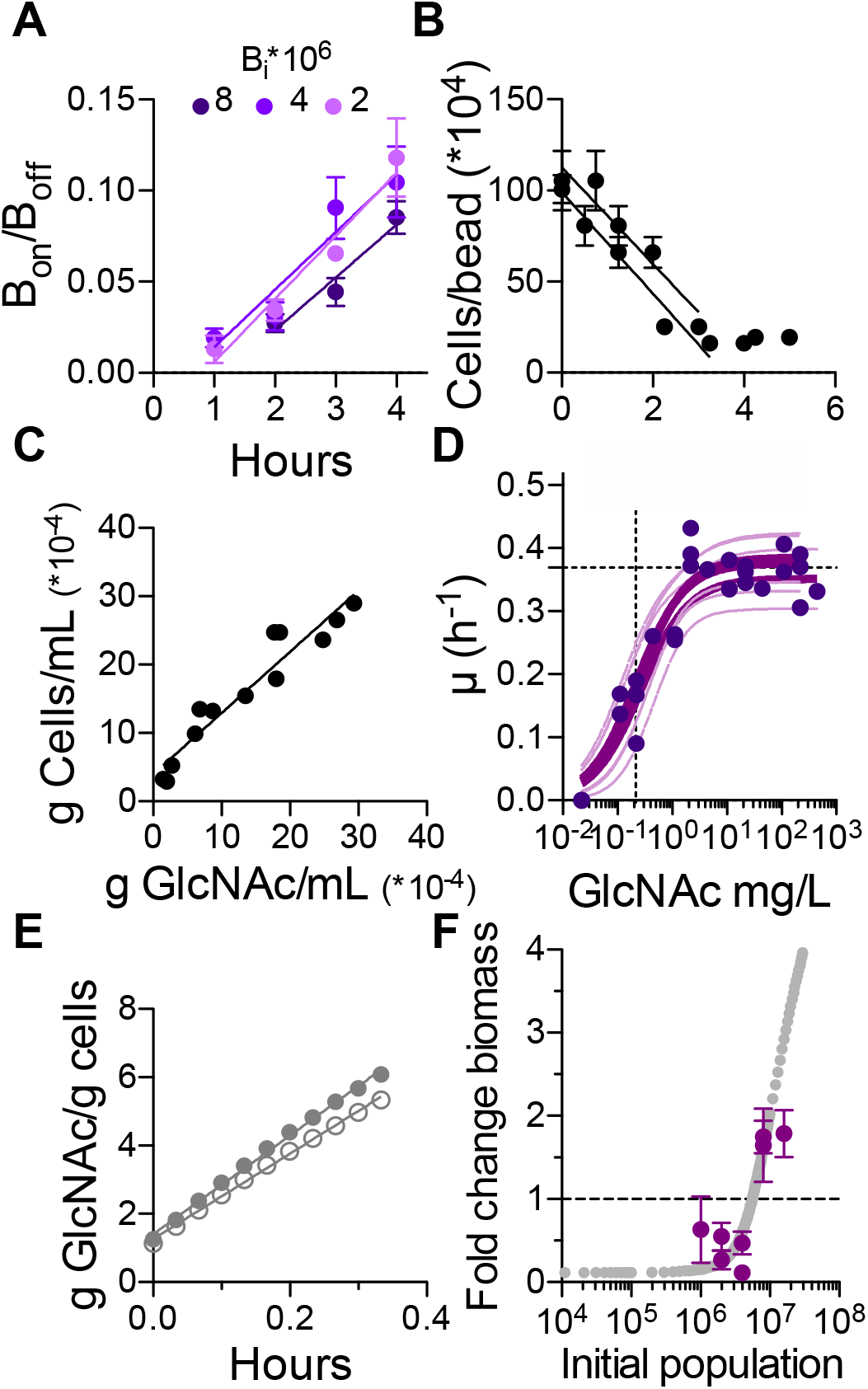
Physiological traits predict an Allee effect for chitin-degrading strain psych6C06 in a diffusive environment. A) Rates of attachment to chitin hydrogel beads, measured for three different initial cell densities per mL (Bi). Error is SEM from a minimum of 10 measurements and lines are fits of the data to a linear regression, where the slope (Bon/Boff) gives the rate of cell accumulation on the particle (for Bi=8*106, Bon/Boff = 2.9*10-2 ± 6.9*10-3; Bi=4*106; for Bon/Boff = 3.2*10-2 ± 6.6*10-3; Bi=2*106, for Bon/Boff = 3.4*10-2 ± 5.0*10-3, ± standard error). B) Rates of detachment from chitin hydrogel beads, measured for two replicates 24 h after colonization by an initial population of 8*106 cells/mL. Error is SEM from a minimum of 11 measurements and lines are fits of the data to a linear regression. Data from the first 3.5 h of the experiment were fit, and the slope of each line gives the rate at which cells leave the particle. Two independent replicates are shown (slope replicate 1= −2.8*105 ± 3.4*104 cells/bead/hour; slope replicate 2= −2.7*105 ± 5.1*104 cells/bead/hour, ± standard error). C) Yield of psych6C06 cells grown on GlcNAc as the sole carbon source. Data points represent individual measurements. Line is a linear regression of the data, where the slope (0.89 ± 0.08 g cells / g GlcNAc) gives the fractional yield. D) Dependency of psych6C06 growth rate on substrate availability. Three biological replicates are shown. Measurements of growth rate at nine different GlcNAc concentrations were fit to the Monod growth model μ = μmax*S/(Ks + S), where μ is the observed exponential growth rate, S is the concentration of GlcNAc, to find the maximum growth rate (μmax) and half-saturation constant (Ks). Bold lines indicates fit for each biological replicate, dashed lines indicate 95% confidence intervals. For replicates 1-3 respectively, μmax=0.35 ± 0.02, 0.39 ± 0.02, 0.37 ± 0.02 and Ks=1.2 ± 0.5, 1.1 ± 0.3, 1.0 ± 0.32. E) The rate of chitin hydrolysis to GlcNAc by cell-associated psych6C06 chitinase (Kp) was measured using fluorescent substrates for exponentially growing cells at two different densities (OD 0.1 open circles and OD 0.4 filled circles). Linear regression was fit to each replicate Kp (OD 0.1)= 12.5 ± 0.2 g GlcNAc/g cells/h, Kp (OD 0.4)= 14.5 ± 0.3 g GlcNAc/g cells/h. The average of the two replicates was used as the estimate for psych6C06 Kp. F) Prediction of psych6C06 biomass yield after 24 h of growth on chitin hydrogel beads with different initial population densities (cells/mL). The analytical model is based on the parameters measured in 3A-E (grey points). Dashed horizonal line indicates no biomass change. Purple data points indicate the experimentally measured change in biomass observed for the indicated initial population densities. Error bars represent SD for measurements from at least 3 replicates.

## Discussion

Despite the key regulatory role of microbes in carbon cycling (17, 26, 27), linking the micro-scale physiology and behavior of bacteria to carbon flux models remains elusive. Here, we show that in conditions where diffusion limits oligosaccharide accumulation, like in the ocean, the breakdown of particulate polysaccharides is subject to population density-dependent effects. These effects are driven by three key physiological parameters: the affinity of cells for hydrolyzed oligosaccharide, the rate of polysaccharide hydrolysis, and the amount of exchange on/off the particle surface that define a tradeoff between the rate of polysaccharide degradation and biomass yield. These features contrast with a common assumption of carbon flux models: that model heterotrophic cells consume nutrients at rates comparable to the consumption of simple dissolved substrates by laboratory-adapted model organisms in a well-mixed systems (27–29). In particular, our computational and experimental results highlight that the frequency at which cells exchange on and off the polysaccharide surface has a large impact on cell carbon uptake. This effect is emerged not only by setting the threshold population density that is achieved by the initial population without growth, but by reconfiguring the arrangement of surface-associated cells in ways that maximize oligosaccharide uptake and growth. In addition, we show that the emergence of microbial aggregates on recalcitrant particles increases the chance of survival for populations of bacteria by enhancing the local dissolved carbon production rate and cell uptake rates. The benefit of aggregate formation is dependent on size: structures that are too large or dense to support the maximum uptake rate of individual cells promote competition rather than cooperation (30). In natural ecosystems, aggregate formation and dispersal is the rule rather than an exception (31–34) and cells are likely to carry adaptations to enhance aggregation/dispersal on particle surfaces, for instance by regulating chemotactic movement or the expression of biofilm components such as adhesins and matrix proteins (35, 36). Thus, micro-scale interactions could significantly affect the rates of POM turnover in the environment, underscoring the need to incorporate these interactions into models of carbon cycling.

Our computational and experimental system also highlights the importance of social collective behavior and spatial self-organization on community fitness and survival. Previous studies have shown that secretion of public goods (enzyme) favors the formation of patchy microbial aggregates by enhancing cooperative behavior (11, 14, 15, 37). Notably, many bacteria actively regulate enzyme secretion and other group behaviors at the level of transcription (38–40), and cell-density-dependent transcription factors such as quorum signal receptors are capable of sensing both changes to the environment and to cell density (41). While our simulations reveal that physiology alone is sufficient to explain patch formation, further studies are required to evaluate the contribution of such regulation on the group behaviors that may facilitate patch formation and dispersal by *psych6C06*. Our results also suggest that the benefit of clustering is not universal and is instead dependent on physiology: the POM uptake efficiency and growth rate of cells with high affinity for substrates suffers in the context of an aggregate, while aggregation is optimal for strains with lower substrate affinity and lower rates of polysaccharide hydrolysis (11). This observation highlights the fact that in systems with potential for spatial organization – that is, most systems outside the lab, the balance between cooperation and competition can be delicate and modulated by the intersection of physical processes with microbial physiology.

## Methods

### Individual based model of individual cell behavior and physiology

The mathematical model represents metabolism, surface interaction and flagellar motility of individual cells in 3D space in the presence of chemical gradients. We introduce an individual-based model (42, 43) to quantify single-cell interactions with organic particles by abstracting the structural heterogeneities of natural POM into a mathematically simpler spherical shape, while preserving some key physical and chemical processes associated with POM degradation. A spherical organic particle of 200 μm radius is simulated such that it remains static in the middle of an aqueous volume (~1mm^3^). While natural organic matter aggregates may show various shapes and chemical compositions, we modeled particles as perfect spheres made of a single type of insoluble linear polysaccharides such as chitin, alginate, or cellulose. This computational model is inspired by experimental model systems used to study community assembly on marine POM (10, 17). The particle’s size and its surface chemistry are assumed to be unchanged during particle degradation: only the particle density changes over time to satisfy mass conservation. This assumption is consistent with experimental observations that have shown no significant change in organic particle size during microbial degradation until the final stages of collapse(17). We simulated a scenario where an isogenic population of cells is allowed to colonize and degrade a particle with a defined volume. The simulations were started with zero oligosaccharides and the particle was considered to be the sole carbon source.

To take into account the fact that cells might regulate their enzymatic activity, the model limits enzyme secretion to two scenarios: when cells adhere to the particle surface or when the rate of oligosaccharide supply exceeds the maintenance threshold. Importantly, our simulations ensure mass conservation between total carbon uptake, growth and loss of oligosaccharides. Individual cells are initialized as a uniform random distribution in the aqueous volume, and are allowed to disperse following gradients of chemo-attractant (in this case, oligosaccharide). The cells can consume the oligosaccharide, grow, and divide to new daughter cells and experience a range of local conditions. A full derivation of the mathematical expressions and steps used for modeling of microbial growth, dispersal and enzyme secretion can be found in the Supplemental Information.

#### Experimental methods

Strain *psych6C06* was previously isolated from an enrichment of nearshore coastal seawater (Nahant, MA, USA) for surface-associated chitin degrading microbial communities (10, 17). The strain was maintained as colonies on Marine Broth 2216 (Difco 279110) with 1.5% agar (BD 214010). To establish exponential growth without hysteresis, we modified a culturing protocol previously developed for *Escherichia coli* K12(44), and grew cells on a defined seawater medium with the *N*-acetyl-_D_-glucosamine (GlcNAc) at concentrations indicated. Chitin hydrogel beads (NEB) were washed and diluted to 200-250 particles per mL with size range from 40 to 100 μm in diameter. The beads were rotated end over end at 21-25 °C. The density of inoculated cells was set to be at an A_600_ of 0.01, diluted from 20 mM GlcNAc minimal medium cultures prepared as described above. To visualize particles and their surface-associated bacteria, 200 μl subsamples were stained with the DNA-intercalating dye SYTO9 (Thermo Fisher, S34854) at a 1:285 dilution of the stock in 96-well plates with optically clear plastic bottoms (VWR 10062-900).

Cell density measurements (Absorbance at 600 nm, A_600_) of exponentially-growing cells were used to measure the maximum cellular growth rate, and plating was used to measure growth under GlcNAc limitation, from which we derived the half-saturation constant. GlcNAc depletion was measured during growth using the dintrosalicylic acid reagent method(45), and the depletion rate was used to calculate the biomass yield (see Supplemental Information)

Chitinase activity was quantified using Methylumbelliferyl(MUF)-conjugated substrates N,N′-diacetyl-β-D-chitobioside, N-acetyl-β-D-glucosaminide, and β-D-N,N′,N″-triacetylchitotriose (Sigma CS1030). Microscopy was performed on micro-confocal high-content imaging system (ImageXpress Micro Confocal, Molecular Devices), using the 60 μm pinhole spinning disk mode. Fluorescent signal was visualized with a LED light cube (Lumencore Spectra X light engine), and bandpass filters (ex 482/35 nm em 538/40 nm dichroic 506 nm), with a 40x objective (Nikon Ph 2 S Plan Fluor ELWD ADM 0.60 NA cc 0-2 mm, correction collar set to 1.1), and a sCMOS detector (Andor Zyla). Image analysis was performed in MATLAB (release 2018a). Briefly, image stacks were split in half and a maximum intensity projection was obtained for each half. The low level of fluorescent signal associated with free dye in the hydrogel particles was used to define an intensity threshold suitable to create a binary mask for the particle projections. A mask of the cells within the beads was then defined using their brighter fluorescence intensity. We used this segmentation to quantify the total surface area occupied by the cells on the bead, and to quantify the total surface area occupied by patches (areas where cells contact other cells >10 μm^2^).

## Acknowledgements

We thank Lu Lu for technical assistance, and all members of the Cordero lab for their support and critical feedback. This project was supported by Simons Early Career Award 410104 and the Simons Collaboration: Principles of Microbial Ecosystems (PriME), award number 542395. A.E. acknowledges funding from Swiss National Science Foundation Grant P2EZP2 175128.

## Supplementary Methods

### Individual based modeling procedure

#### Growth and division

The individual-based model assumes that an individual cell doubles to two identical cells after a certain amount of carbon is taken up (1, 2). In this model, a Monod-kinetic parameterization(3) gives the carbon uptake kinetics of individual cells 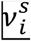 and their biomass accumulation:

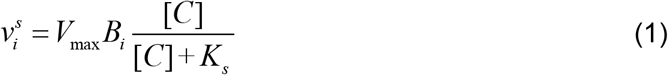

where *B*_*i*_ is the cell dry mass of the individual cell, *i*, and the maximum rate of substrate uptake is: *V*_max_=*μ*_max_ / *Y*_max_ (maximum specific growth rate / growth yield). *K*_*s*_ is the half-saturation constant for dissolved oligosaccharide. We assume that dissolved oligosaccharides [*C*] are the primary limiting substrate for growth, and that all other nutrients (e.g., sources of phosphate and nitrogen) available at non-rate limiting levels.

For cell *i*, the actual biomass accumulation 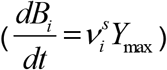 and resources consumed (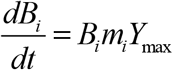) for maintenance, *m*_*i*_ are both assumed to be propositional to the cell’s dry mass, *B*_*i*_, therefore, the net growth in the cell biomass (*μ*_*net*_) can be described as follows:

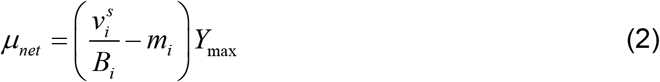

The minimum volume of the individual cell at division (*V*_*d, min*_) is estimated from the descriptive Donachie model (1, 2)

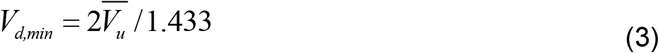

where 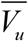 is median volume of the individual cell. If the individual cell volume becomes larger than *V*_*d, min*_, the parent cell divides to form two new daughter cells. The individual cells are assumed to be cylindrical. It should be noted that the actual growth kinetics of an individual cell could be limited due to substrate availability within the corresponding mathematical mesh grid (*M*_*s*_) at each time step (Δt), therefore if the uptake driven by Eq. 1 is higher than the available substrate in a given mesh grid 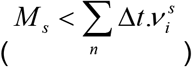 then the available substrate is equally shared among the number of individuals inhabiting the same grid (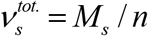, n: number of individuals within the same mesh grid). The biological parameters of the kinetic microbial model are summarized in Table S2.

#### Microbial active movement

Microbial movement is modeled as flagellar motility: a self-propulsive force guided by chemotaxis that is driven by local substrate gradients (4, 5). The model describes the chemotactic movement of an individual bacterium as a biased-random walk where the flagella propel a straight “run” by rotational movement of the motor and then reverse the rotation to switch direction with a probability to change direction (“tumble”) set by surrounding gradients of chemo-attractants. To apply chemotactic movement to a single cells at each time step, the sensitivity of microbial chemoreceptors towards higher substrate concentrations are described using a receptor model that uses the specific growth rate as the chemotactic potential (6, 7). Thus in the biased random walk of an individual bacterium, the probability of transition *P*_*move,i*_ in a tumbling event into a new (more favorable) direction from the current direction (previous “run”) is expressed quantitatively as(1, 6):

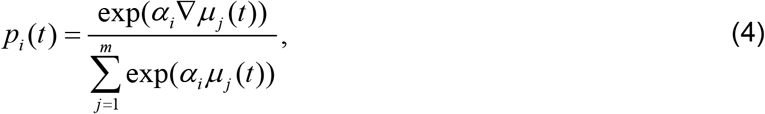

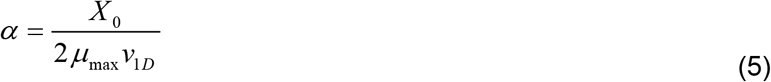

Where α is the factor for the chemotactic motion derived from chemotactic sensitivity coefficient of individual cells to chemo-attractant(8), *X*_0_, bacterial maximum swimming velocity, *ν*_1*D*_ and the growth rate in response to local gradients, *μ*_*max*_. *m* is the number of directions that an individual cell could sense the local gradients in 3D surrounding environment. To insure homogenous directional choices, the summation is evaluated with a relatively high number of possible directions (*m*=20) for each individual cells, but still minimize the computational burden when scaling up to systems of thousands of individual cells. The distance in which bacterial cells sense the local chemotactic gradient is set equal to the cell length (1 μm). Periodic boundary conditions are assumed for bacterial cells that pass outside of the external boundaries of the system.

#### Bacteria-particle interactions

Bacterial cells are allowed to attach to the particle surface when they stochastically encounter a particle. The model assumes that bacterial cells attach to the particle upon surface encounter, but that a set detachment probability for each cell allows them to detach from the particle. This probability is invariant over the span of time that cells are surface-associated, which contrasts with many characterized mechanisms by which bacteria form irreversible contact with surfaces(9). To determine the effect of this simplifying assumption on the behavior of the model, we simulated a wide range of detachment probabilities, ranging from no detachment to a relatively high detachment rate (1% detachment probability per second).

#### Enzymatic activity

Cells in the model can broadcast extracellular enzymes. These enzymes are produced at a constant rate that is proportional to the biomass of the cell, and the enzymes can diffuse into the bulk environment. Enzymes that come into contact with the particle subsequently hydrolyze polysaccharide and release diffusible oligosaccharide at the particle surface. The production of enzyme is assumed to be activated when a bacterial cell adheres to the particle, or when cells take up enough oligosaccharide to support cell maintenance. That is, the process of enzyme production preserves mass conservation. We modeled the rate that an individual cell *I* broadcasts enzymes (*S*_*E, i*_) as a conditional linear function to its biomass, *B*_*i*_:

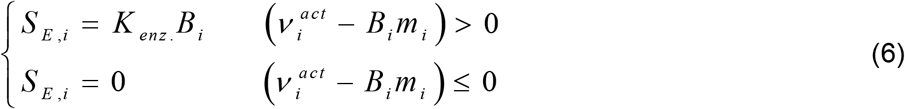

where 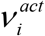 is the actual uptake rate by the individual cell and *m* is the maintenance rate. The diffusion of the enzyme is then solved by considering a source term, equivalent to the total production rate of enzymes from a corresponding cubic mesh grid 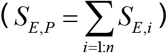 and a sink term on the particle source that gives the rate at which enzymes adhere (*S*_*A, P*_).

#### Oligosaccharide diffusion-reaction in physical domain

The model explicitly simulates the loss of oligosaccharides to the bulk environment due to the presence of diffusion or flow. This is implemented by absorbing conditions at the boundaries of the simulated domain in which oligosaccharides that arrive at a boundary are lost to the bulk environment with no accumulation. This is a relevant assumption for many aquatic and terrestrial ecosystems, in contrast to soil ecosystems under dry conditions that may impose restrictions on substrate diffusivity or to batch culture where substrates accumulate in culture vessels.

The transport and uptake of depolymerized oligosaccharides are modelled based on Fick’s law of diffusion and mass conservation. We modeled a 3D cubic volume around a single particle and assumed that diffusion of oligosaccharides is the main mode of mass transport. The reaction-diffusion equation is then numerically solved by the finite-difference method, assuming a regular cubic mesh discretization(10):

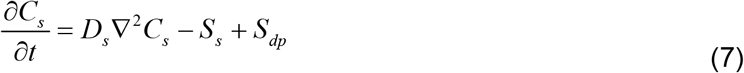

where *C*_*s*_ is the concentration of oligosaccharide (the product of enzymatic hydrolysis), *D*_*s*_ is the diffusion coefficient of the oligosaccharide and *S*_*s*_ is the oligosaccharide consumption rate (the sink term) due to microbial uptake. S_s_ is a summation of the individual uptake rates (*ν*^*s*^), of all cells within a cubic mesh grid (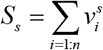, n is the number of individuals in the mesh grid) (10). *S*_*dp*_ is the production rate of oligosaccharide given from Eq. 11. The mesh grid size is chosen to be approximately 10 μm and the time step used for simulating oligosaccharide diffusion is the same as that used for chemotactic movement for computational simplicity. For each time step, the Dirichlet boundary conditions are applied for the particle surface and the external boundaries of the cubic volume around the particle. The concentration of oligosaccharide at the external boundaries is set to zero to create an absorbing boundary condition (eliminate the accumulation of oligosaccharide). This external boundary condition is similar to that used for enzyme diffusion (below). A convective term is added in the case where oligosaccharide transport processes are modeled around sinking particles.

#### Enzyme diffusion and decay

The diffusion of the enzyme is solved by considering a source and sink term, similar to Eq. 7:

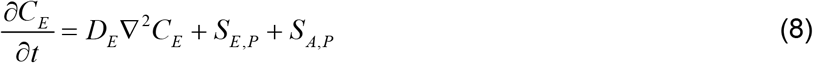

here *C*_*E*_ is the concentration of broadcast enzyme and *D*_*E*_ is the enzyme diffusion coefficient. Both values can be measured experimentally. In this study, we used an empirical model that decomposes the diffusion coefficient into components describing the viscosity *η* and temperature *T* of the medium, and molecular weight of the molecule, *Mw D*_*E*_ = 1.7×10^−7^*T*/*Mw*^0.41^*η* (cm^2^/s) (11). In Eq. 7, the adhesion of enzyme to the particle acts as a sink term at the particle surface boundary that is equivalent to the total amount of enzyme that arrives at the particle surface (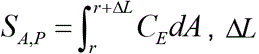). The total enzyme accumulation on the particle (*S*_*E,surf*_.) is then the integral of the accumulation rate *S*_*A,P*_ minus the decay rate, *S*_*E,D*_:

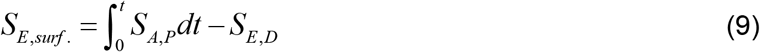

Enzymes are assumed to decay with first order kinetics, so *S*_*E,D*_ is a function of the amount of enzyme adsorbed to the particle surface (12–14):

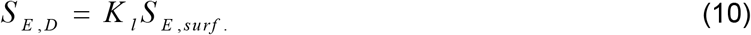

where *K*_*l*_ is defined as the enzyme decay coefficient. The depolymerization rate of polysaccharide (*S*_*dp*_) to oligosaccharide is therefore a function of the particle-adsorbed enzyme (*S*_*E,surf.*_) with a linear empirical relationship:

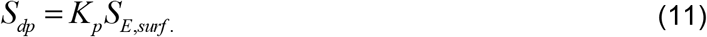

where *K* _*p*_ is the particle lability, defined as a lumped parameter that resembles biopolymer and enzyme biochemistries. That is, *K* _*p*_ is a combined term to express difficulty of the particle to degrade relative to the activity of the enzyme produced by the cells.

#### Oligosaccharide transport in presence of particle sinking

Where we model sinking in our simulations, we assume a constant 1D speed rate along the water column, *u* (see Figure S2). For simplicity, we assume that particle sinking creates a laminar flow around the particle with the same speed as the particle sinking rate. To address the effect of flow on dissolved carbon and enzyme transport, we added advection term to diffusion model for dissolved carbon and enzyme transports (Eq. 7 and 8) and expressed as:

For dissolved carbon:

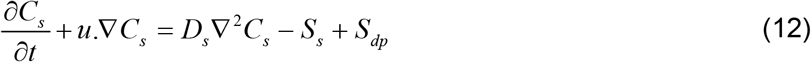

For enzyme:

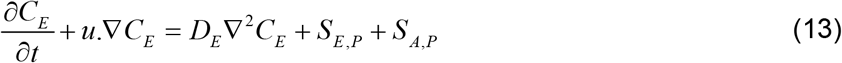

where sink and source terms for dissolved carbon (*S*_*s*_, *S*_*dp*_) and enzyme (*S*_*s*_, *S*_*dp*_) are all expressed similar to equations 7 and 8, respectively. Similar to the stationary particle simulations, the mesh grid size was chosen to be approximately 10 μm and the time step of calculations was assumed to be similar to the time interval between chemotactic tumbles (5 seconds). At each time step, Dirichlet boundary conditions are applied to the particle surface to the external boundaries of the volume around the particle. At the particle surface a constant flux of oligosaccharide production is considered as given by Eq. 11, as a function of the particle-adsorbed enzyme concentration. As above, the external boundary concentration of oligosaccharide is set to zero. A finite difference method was applied to solve Eq. 12 and 13 simultaneously at each time step within the individual-based model.

To model longer time scales of particle sinking in the water column, we implemented a spatial algorithm to only model the effective zone around the particle in each time scale, instead of modeling the whole water column for each time step. Based on analyzing the concentration gradient of dissolved carbon around the particle, we chose the effective zone as the region with concentration gradient above a threshold value (5% of maximum chemical gradient).

### ‘Population-level’ analytical bottom-up model

We developed a simple quantitative model to predict the fold change in biomass based on the measurable physiological and behavioral features of marine bacterial isolates. The model is parametrized based on the rate of enzyme production and activity (degradation rate) to predict the total of oligosaccharide production, *M* over time, given by a first order kinetics:

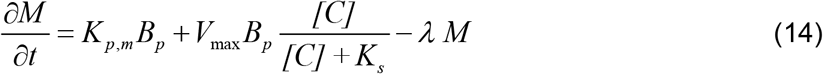

where *K*_*p,m*_ is the rate of oligosaccharide production per biomass per hour that is derived from enzymes bound to the cell membrane. *B*_*p*_ is the particle-associated fraction of the total biomass. 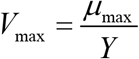 is the maximum uptake rate, defined as the ratio of maximum growth rate of bacteria, *μ*_*max*_ to the yield of substrate conversion to biomass, *Y* (experimentally measured). *λ* is the fraction of monomers that are lost to the bulk environment and the value is assumed from what we obtained from simulations individual based model, and from previous reports in the literature about the inefficiency of oligosaccharide recovery from hydrolysis(15). Note that as the enzyme is tethered to the cells, no diffusion for the enzyme is assumed.

The particle-associated biomass production rate is represented based on the combination of Monod-type growth kinetics and attachment-detachment frequencies:

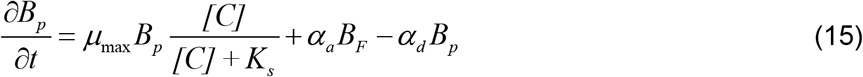

where *B*_*F*_ is the fraction of free living biomass in the system, *[C]* is the oligosaccharide concentration, calculated from the ratio of oligosaccharide mass, *M* to the volume. *K*_*s*_ is the half-saturation constant, experimentally measured in Figure 4. *α*_*a*_ and *α*_*d*_ are attachment and detachment rates, respectively, measured in Figure 4.

The change in free-living biomass, *B*_*F*_ is derived from the frequency of attachment and detachment, assuming that free-living cells do not themselves grow:

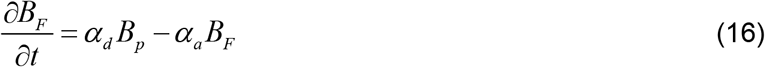

### Experimental methods

#### Culture conditions

Strain *psych6C06* was previously isolated from an enrichment of nearshore coastal seawater (Nahant, MA, USA) for surface-associated chitin degrading microbial communities (16, 17). The strain was maintained as colonies on Marine Broth 2216 (Difco 279110) with 1.5% agar (BD 214010). To establish exponential growth without hysteresis, we modified a culturing protocol previously developed for *Escherichia coli* K12(18). Briefly, single colonies were picked and transferred to 3 mL liquid Marine Broth 2216 and incubated at 25 °C, shaking at 150 rpm on a VWR DS-500 orbital shaker to establish seed cultures. Seed cultures were harvested after ~5 hours by centrifugation for 1 min at 5000 rcf (Eppendorf 5415D, Rotor F45-24-11). The supernatant was discarded and serial dilution of the cells were used to establish pre-cultures in pH 8.2 minimal media supplemented with 20 mM *N*-acetylglucosamine (GlcNAc). The core minimal medium contained the major ions present in seawater, plus vitamins and trace minerals per L: 20 g NaCl, 3 g MgCl_2_-6H_2_O, 0.15 g CaCl_2_-2H_2_O, 0.05 g KCl, 2.1 mg FeSO_4_-7H_2_O, 30 μg H_3_BO_3_, 100 μg MnCl_2_-4H_2_O, 190 μg CoCl_2_-6H_2_O, 2.2 μg NiSO_4_-6H_2_O, 2.7 μg CuSO_4_, 144 μg ZnSO_4_-7H_2_O, 36 μg Na_2_MoO_4_-2H_2_O, 25 μg NaVO_3_, 25 μg NaWO_4_-2H_2_O, 2.5 μg SeO_2_, 100 μg riboflavin, 30 μg D-biotin, 100 μg thiamine-HCl, 100 μg L-ascorbic acid, 100 μg Ca-_D_-pantothenate, 100 μg folate, 100 μg nicotinate, 100 μg 4-aminobenzoic acid, 100 μg pyridoxine HCl, 100 μg lipoic acid, 100 μg nicotinamide adenine dinucleotide, 100 μg thiamine pyrophosphate, and 10 μg cyanocobalamin. In addition to the carbonate buffer present in the core minimal medium (reflecting natural seawater buffering capacity), we added 50 mM HEPES buffer pH 8.2 to control for the effects of heterotrophic metabolism on pH. We also supplemented the core medium with10 mM NH4Cl, 1 mM Na_2_PO_4_, and 1 mM Na_2_SO_4_ to create a carbon-limited minimal medium. Where appropriate, we supplemented with the *N*-acetyl-_D_-glucosamine (GlcNAc, concentrations indicated). Following overnight growth at 25 ºC, the cell density was measured in 1 cm cuvettes by absorbance measurements at 600 nm (A_600_) using a Genesys 20 spectrophotometer. Under these conditions, A_600_ 1.0 =8×10^8^ cells/mL, measured by serial dilution and plating.

### Measurement of Monod growth parameters

To measure the maximum growth rate on *N*-acetylglucosamine, cells were prepared as described above, then diluted to a starting OD of 0.01 in 4 mL of fresh 20 mM GlcNAc minimal medium in a 10 mL vented polystyrene tube. OD measurements were taken over time, and the linear regression was fit to the plot of ln(OD) vs. time. The slope of this regression is equivalent to the growth rate. To measure the half-saturation constant, cells were diluted from a OD 1.0 by a factor of 10^5^ into carbon-free minimal medium, and 100 μl of this dilution was used to inoculate medium containing a GlcNAc at different concentrations (221, 110, 22, 11, 2.2, 1.1, 0.22, 0.11, and 0.02 mg/L). At 2 h time intervals, 100 μl of this culture was plated onto MB 1.5% agar plates. The resultant colonies were counted, and the change in colony numbers over time was used to derive the growth rate. The maximum growth rate measured by this method was the same as that measured using the OD-based approach. Growth rate was plotted against carbon concentration (S), and the experimentally measured μ and S were fit to the Monod growth equation (μ=μmax(S/(S+K_sat_)) to derive parameters μmax and K_sat_ using a least squares fit with a maximum of 1000 iterations. The biomass yield during growth on GlcNAc was derived from direct measurement of cell density (A_600_) and GlcNAc depletion using the dintrosalicylic acid reagent method to colorimetrically quantify reducing sugars in cell-free media(19). The grams of GlcNAc/mL depleted from the media was plotted against the grams of cells produced, at timepoints covering three population doublings (A_600_ 0.1−0.8). The mass of an individual cell was assumed to be 19 fg: a value which we derived from scaling the measured mass of individual E. coli with 60 min doubling time (220 fg)(20), scaled by the growth rate of *psych6C06* (0.35), and also by a factor of 4 to reflect a linear increase in biomass per cell with osmolarity between M9 and seawater (21), divided by the ratio of *E. coli* to *psych6C06* volume (16:1).

### Measurement of protein production and enzymatic activity

To collect secreted protein and cell-associated protein, cells were grown in large batches. Cells were prepared for inocualtion as described above, and inoculated at an initial density of OD 0.01 into 250 mL Erlenmeyer flasks containing 150 mL of 20 mM GlcNAc minimal medium. The flasks were grown with 150 rpm shaking at 25 °C. Periodically, 25 mL of culture was removed from the flasks, and centrifuged at 5000 rcf for 10 min at 4 °C. Sampling was stopped when the culture flask volume reached 75 mL, past which point culture growth rate was affected by volume. The supernatant was collected and filtered through 0.2 μm Sterivex filters (EMD Millipore), at which point protease inhibitor (Roche cOMplete) was added. The pellet was immediately frozen at −20 until quantification. Supernatant was concentrated ~50 x in Amicon Ultra 3 kDa centricon tubes (EMD Millipore), and rinsed twice with 12 mL of carbon-free minimal media to remove small molecules and other potentially inhibitory compounds. The protein abundance was quantified by measuring absorbance at 280 nm, with a 340 nm pathlength correction on a Nanodrop spectrophotometer (Thermo Scientific). The quantification was calibrated using standards of from proteins of known concentration (BSA, chitinase). Chitinase activity was quantified using Methylumbelliferyl(MUF)-conjugated substrates N,N′-diacetyl-β-D-chitobioside, N-acetyl-β-D-glucosaminide, and β-D-N,N′,N″-triacetylchitotriose (Sigma CS1030). Briefly, 2.5 μl of a 20 μg/mL stock of each substrate in DMSO was added to 197.25 μl of concentrated protein or crude cell lysate in chitinase assay buffer (carbon-free minimal medium, pH 8.2 with no vitamins, ammonium or phosphate). The amount of cell lysate was normalized prior to assay. The accumulation of fluorescence (ex 360-20/em 450-20) was monitored on a Tecan Spark at 25 °C with a cooling module, by measuring fluorescence signal accumulation at 2-minute intervals with continuous shaking at 54 rpm with 6 mm amplitude. Serial dilutions of an unconjugated 4-Methylumbelliferone standard were used to establish the linear range of the instrument and to convert fluorescence intensity into mg/mL of released oligosaccharide.

### Bacterial colonization on particles

Precultures with absorbance between 0.1-0.3 were prepared as described above and used to colonize magnetic chitin hydrogel beads. To prepare the beads, 500 μl of bead slurry was washed 3 times with carbon-free minimal media using magnetic pulldown. The washed beads were further diluted 1:3 and used to fill 15 mL conical tubes. The particle density in the tubes was counted, and all replicates contained between 200-250 particles per mL with size range from 40 to 100 μm in diameter. This density of beads is consistent with previous studies of community assembly(17), and provides the equivalent of about 100 μM of the monomeric unit of chitin, *N*-acetylglucosamine (GlcNAc) to the system. The beads were rotated end over end, so that they fell through the medium due to gravity and remained constantly suspended. Because of the rotation, we were unable to continuously observe the beads and instead sub-sampled the population and made individual measurements of multiple beads at each sampled timepoint. The density of inoculated cells was set to be at an A_600_ of 0.01, diluted from 20 mM GlcNAc minimal medium cultures prepared as described above. A vertical wheel (Stuart S3B, 10” diameter wheel) was used to rotate the 15 mL tubes at 5 rpm at room temperature (21-25 °C) with overhead rotation. To visualize particles and their surface-associated bacteria, 200 μl subsamples were stained with the DNA-intercalating dye SYTO9 (Thermo Fisher, S34854) at a 1:285 dilution of the stock in 96-well plates with optically clear plastic bottoms (VWR 10062-900). To avoid evaporation from the wells, sterile self-adhesive sealing films were used to seal the 96-well plates.

### Confocal microscopy and image processing

Microscopy was performed on micro-confocal high-content imaging system (ImageXpress Micro Confocal, Molecular Devices), using the 60 μm pinhole spinning disk mode. Fluorescent signal was visualized with a LED light cube (Lumencore Spectra X light engine), and bandpass filters (ex 482/35 nm em 538/40 nm dichroic 506 nm), with a 40x objective (Nikon Ph 2 S Plan Fluor ELWD ADM 0.60 NA cc 0-2 mm, correction collar set to 1.1), and a sCMOS detector (Andor Zyla). To visualize individual particles, particles were manually centered in the field of view and then 100 μm image stacks sampled at Nyquist were acquired in the Z plane using MetaXpress software (version revision 31201). Image analysis was performed in MATLAB (release 2018a). Briefly, image stacks were split in half and a maximum intensity projection was obtained for each half. The low level of fluorescent signal associated with free dye in the hydrogel particles was used to define an intensity threshold suitable to create a binary mask for the particle projections. A mask of the cells within the beads was then defined using their brighter fluorescence intensity. We used this segmentation to quantify the total surface area occupied by the cells on the bead, and to quantify the total surface area occupied by patches (areas where cells contact other cells >10 μm^2^).

**Figure S1.**
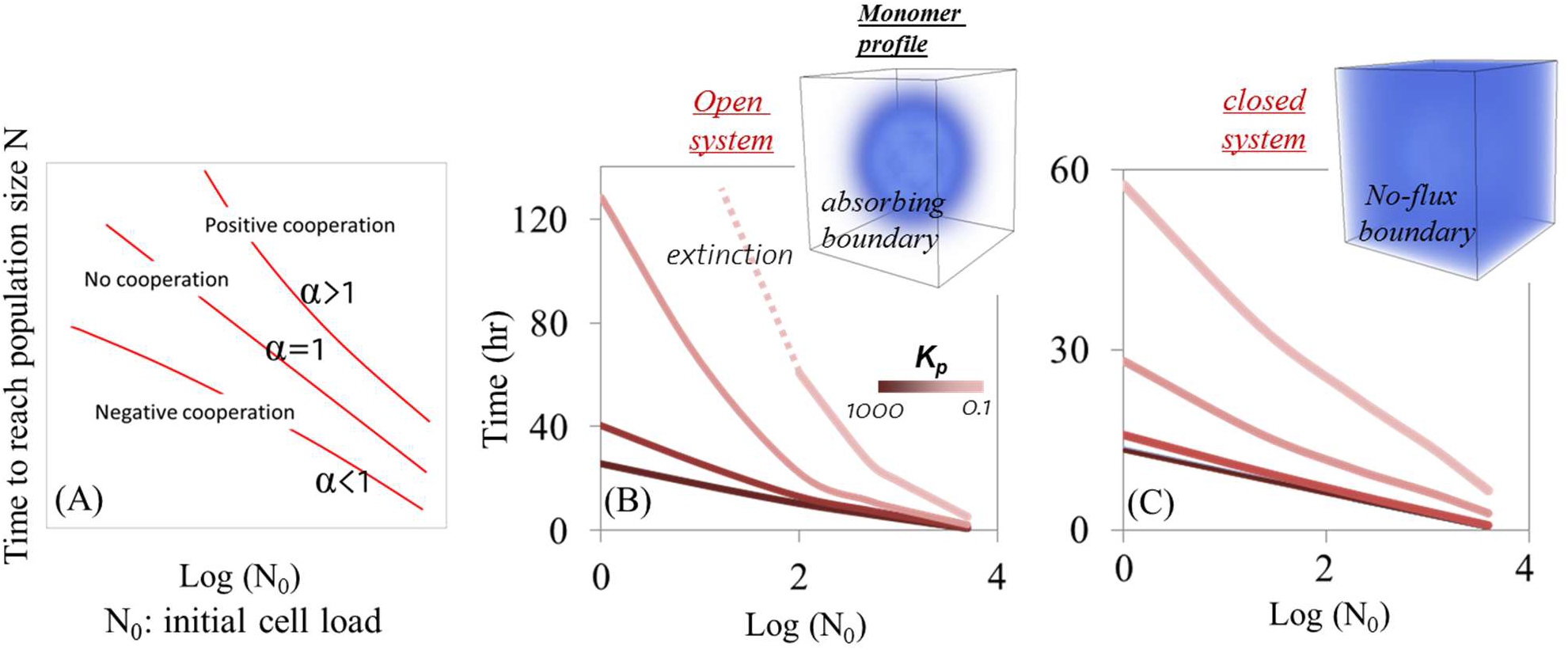
Microbial cooperative behavior affected by batch (closed system with no flux boundary conditions for bacterial cells, dissolved carbon and enzymes) vs. open (absorbing boundary conditions for dissolved carbon and enzyme and periodic boundary for bacterial cells) systems.(A) Cooperative behavior is described as dependency of bacterial growth rate to cell initial load (N_0_). It is defined based on divergence from exponential growth behavior 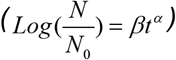 in which is α is 1 for exponential growth and α > 1 shows positive cell dependency of growth rate (Cooperative growth). (B) and (C) growth rate dependency to initial cell load for open and closed systems, respectively. Dashed line indicates the extinction of bacterial community for cell load lower than a threshold value. Inset images indicate the dissolved carbon concentration profiles for their corresponding scenarios. Simulations are done for a particle size of 200μm and results are plotted for the time that allows reaching to population size of 10^5^ cells. Our simulations reveal that diffusion in an open system may lead to the emergence of spatial self-organization and cooperative growth among individual cells (defined as positive dependency of growth rate to cell number). However, in a closed environment, like a batch culture environment (often the case for experimental designs) this need not be the case. With limited diffusion, substrates produced from particulate biopolymers could eventually accumulate and be fully consumed (assuming that no other factors limit growth). Such closed conditions lead to a stepwise relationship between POM uptake efficiency and particle lability where above a threshold lability (K_p_), POM uptake efficiency reaches to 100% and below that no degradation/uptake is expected.

**Figure S2.**
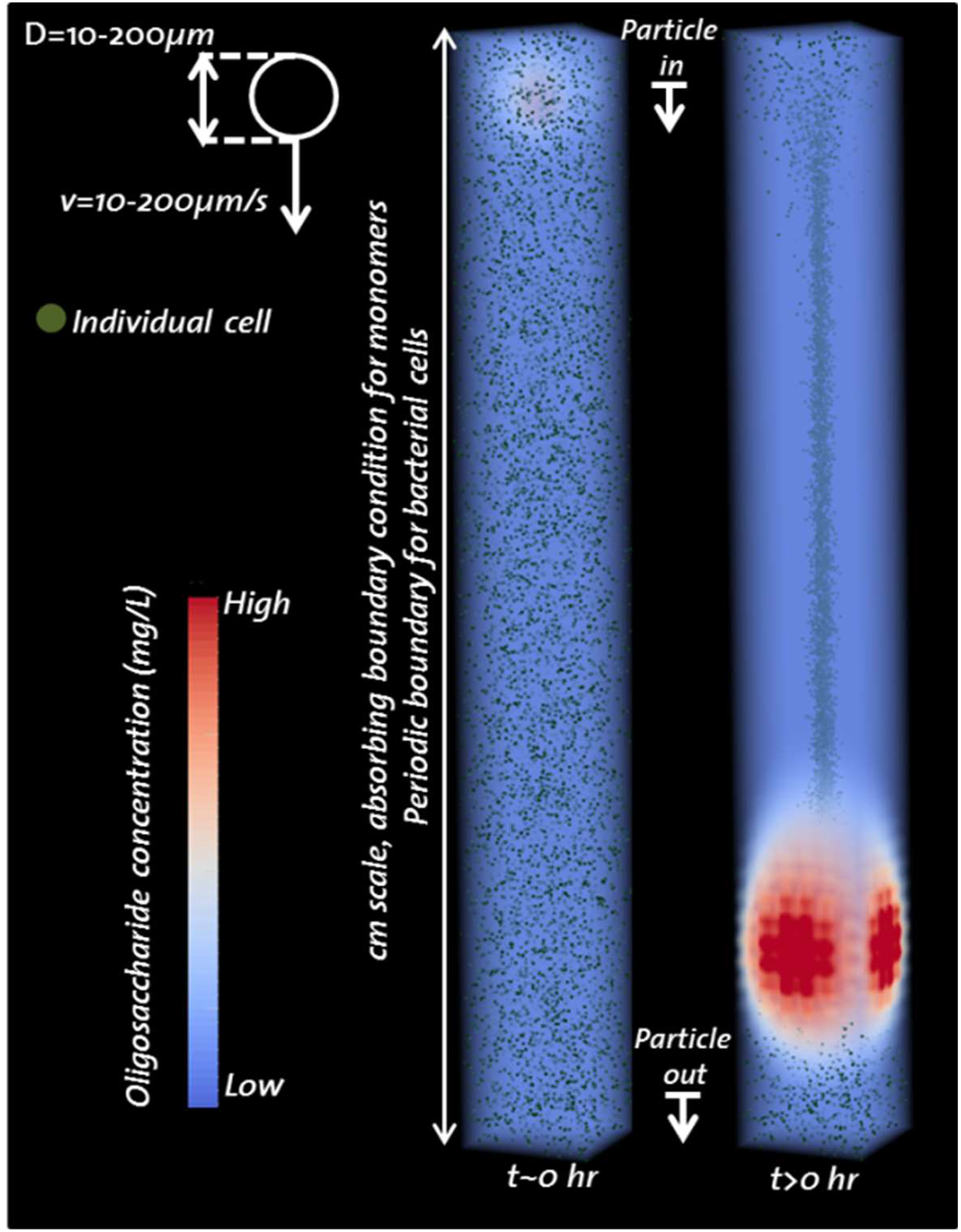
Model description for particle sinking, oligosaccharide profile and individual cell distributions. Individual cells are uniformly distributed over the water column. Each individual is assigned dispersal and enzymatic functions.

**Figure S3.**
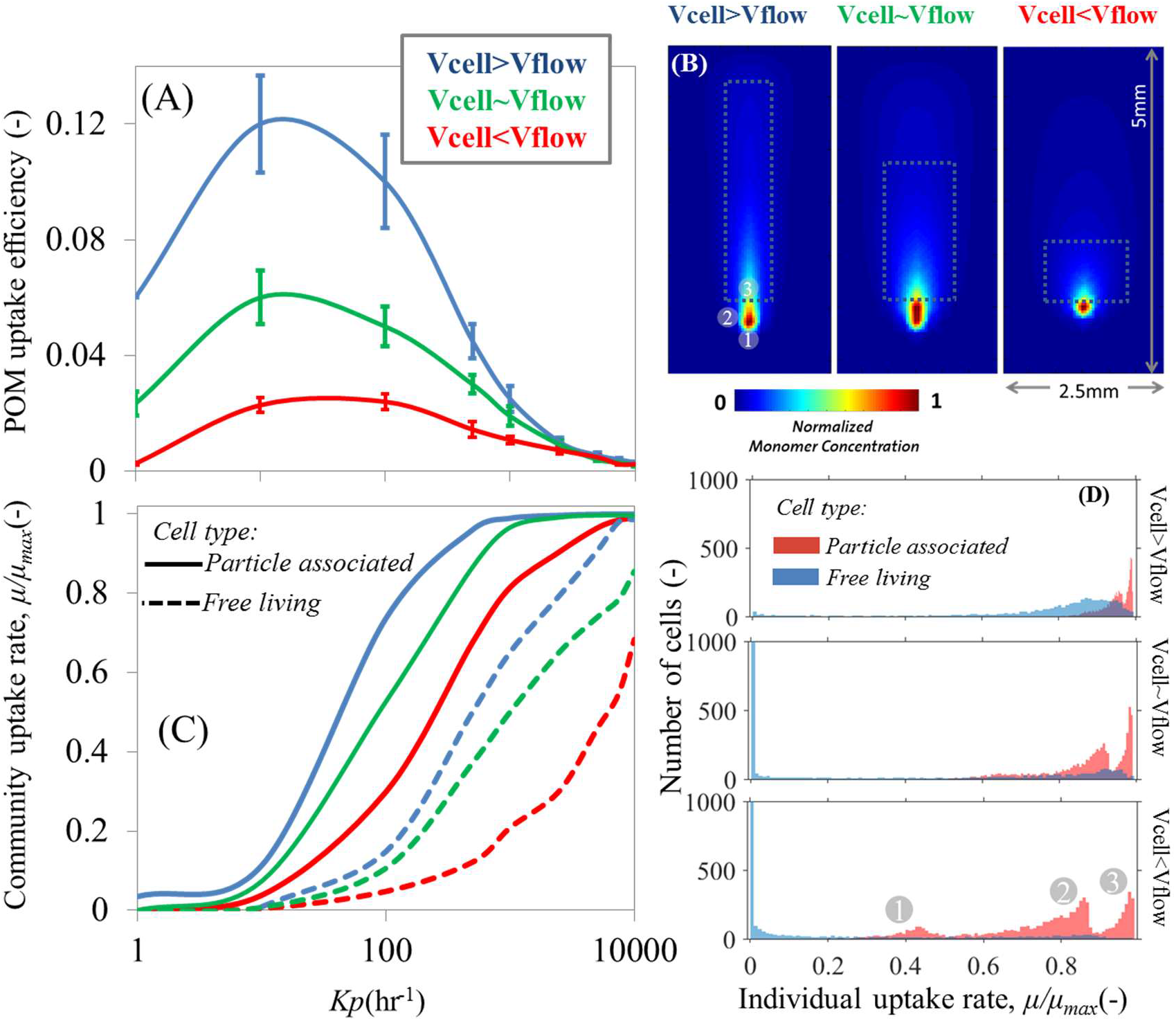
Particle sinking affects local gradients of dissolved organic matter and POM uptake efficiency of particle associated and free-living bacterial cells. (A) POM uptake efficiency for a wide range of particle lability affected by the ratio of cell velocity (Vcell) to particle sinking speed (Vflow). (B) Spatial patterns of dissolved carbon concentration around the particle. Concentrations are normalized by the maximum local concentrations for each scenario. (C) Mean uptake rate at community level presented for particle associated and free-living bacterial cells. (D) Individual cell uptake rate frequencies. The cell velocity is kept constant at 10μm/s and particle sinking speed is changed from 5 (top) to 10(middle) and 20(bottom) μm/s to evaluate the effects of various ratios. The numbers in grey circle at the peaks of histogram for scenario Vcell <Vflow corresponds to the individual cells spatial location around the particle in Figure S8B. The results are shown for simulations after 10hours. Initial cell number is about 10^5^ cells per ml uniformly distributed along the depth of the water column.

**Figure S4.**
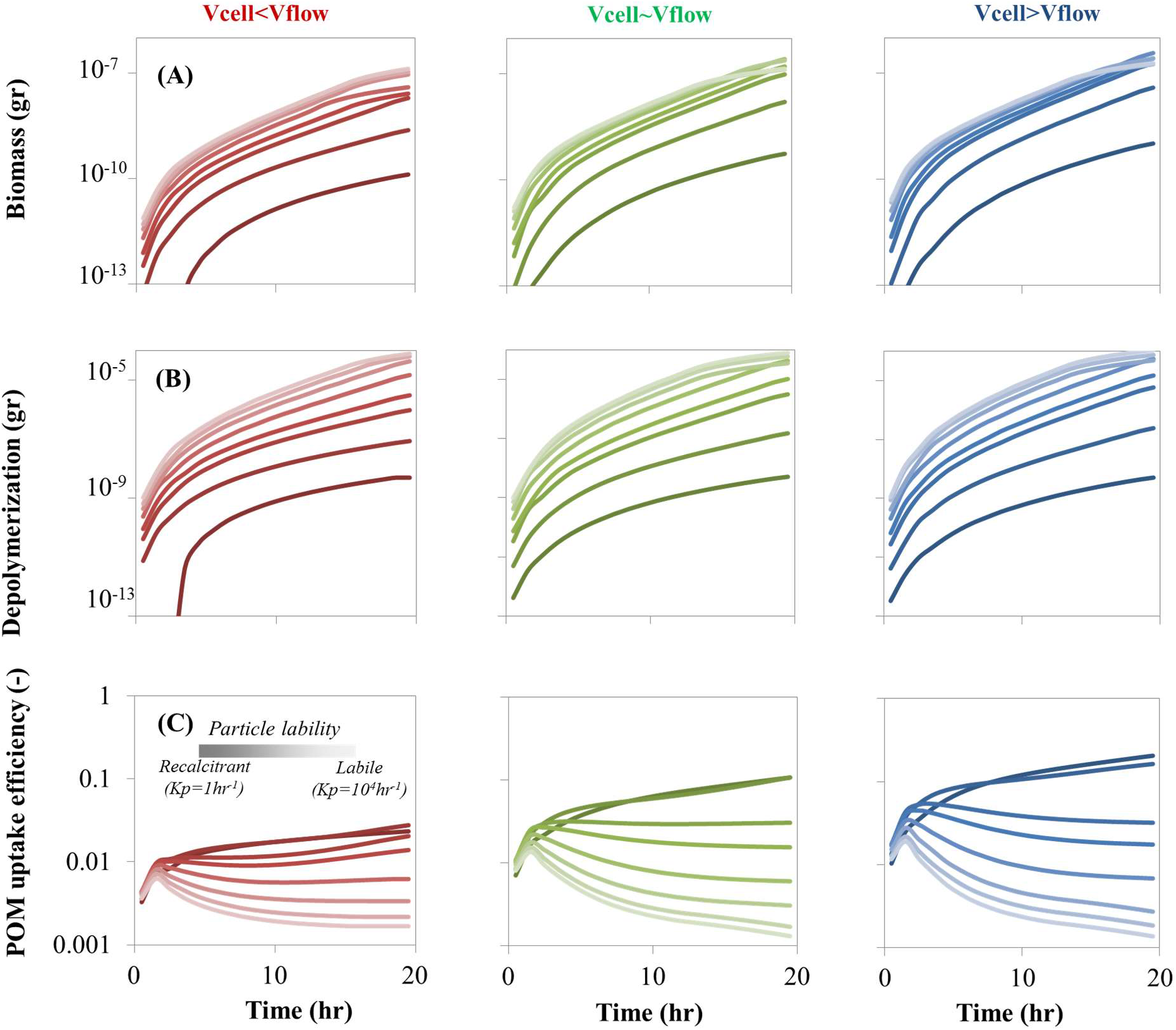
Particle sinking affects biomass accumulation (A) and depolymerization (B) rates and uptake efficiency (C). The simulations are performed for 3 ratios of cell to particle sinking speeds. The results are represented for various polymer labilities from labile (Kp=10000 hr^−1^) to recalcitrant (Kp=1 hr^−1^). The cell velocity is kept constant at 10μm/s and particle sinking speed is changed from 5 to 10 and 20 μm/s to evaluate the effects of various ratios. The results are shown for simulations after 10hours from initializing the simulations. Initial cell number is ~10^5^ cells per milliliter volume.

**Figure S5.**
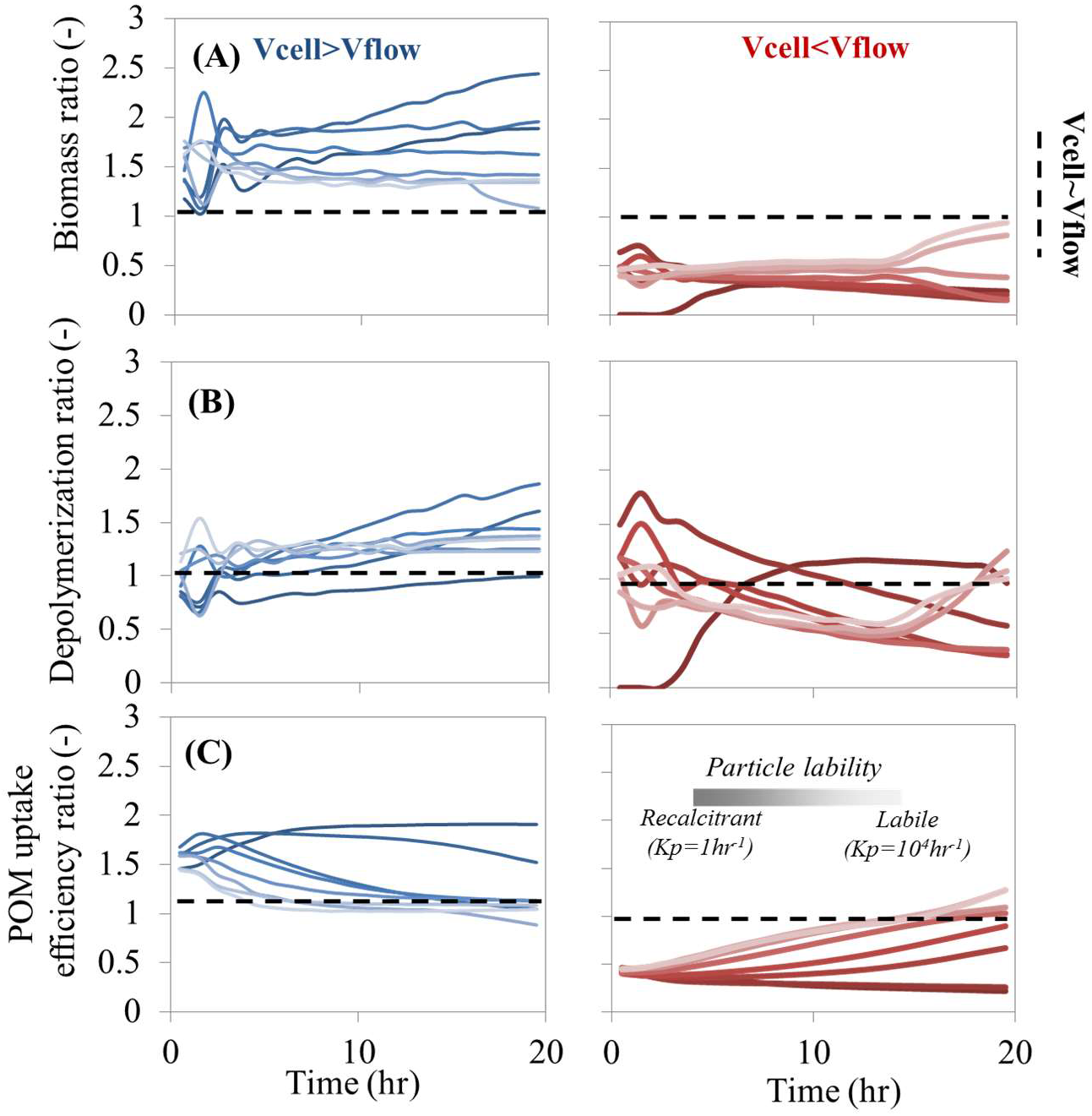
Comparing the effects of particle sinking on biomass accumulation (A) and depolymerization (B) rates and uptake efficiency (C) against reference sinking speed. The reference scenario is assumed to be when cell velocity and particle sinking speed are equal (V_cell_~V_flow_). Ratios are calculated by dividing the biomass accumulation, depolymerization and POM uptake efficiency from Figure S9 for two scenarios of particle sinking (V_cel_ >V_flow_ & V_cel_ <V_flow_) with reference scenario.

**Figure S6.**
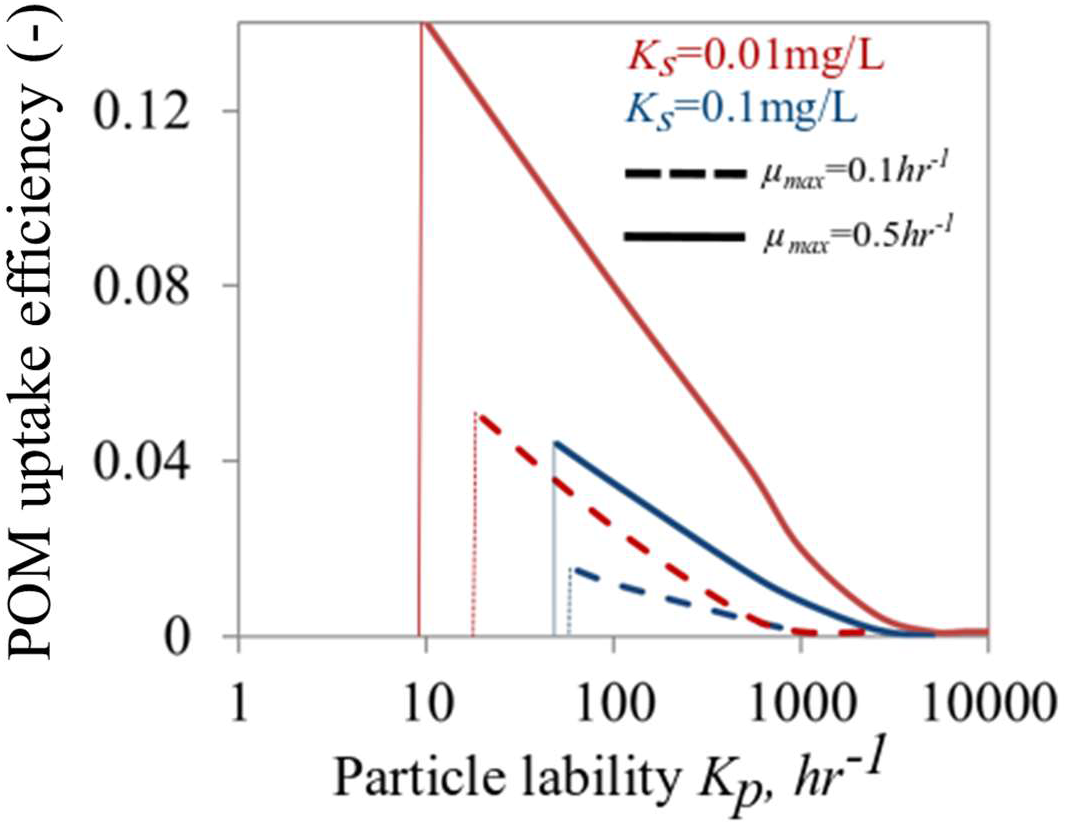
Physiological properties of bacterial cells (maximum growth rate μ, affinity to substrate Ks) affect POM uptake efficiency. POM uptake efficiency as a function of particle lability is shown for high (μ=0.5hr^−1^) and low (μ=0.5hr^−1^) maximum growth rates. The results are shown for two levels of affinity to substrate (0.1 and 0.01 mg/L). Vertical lines show the threshold particle lability below which no particle degradation and growth/substrate uptake is observed.

**Figure S7.**
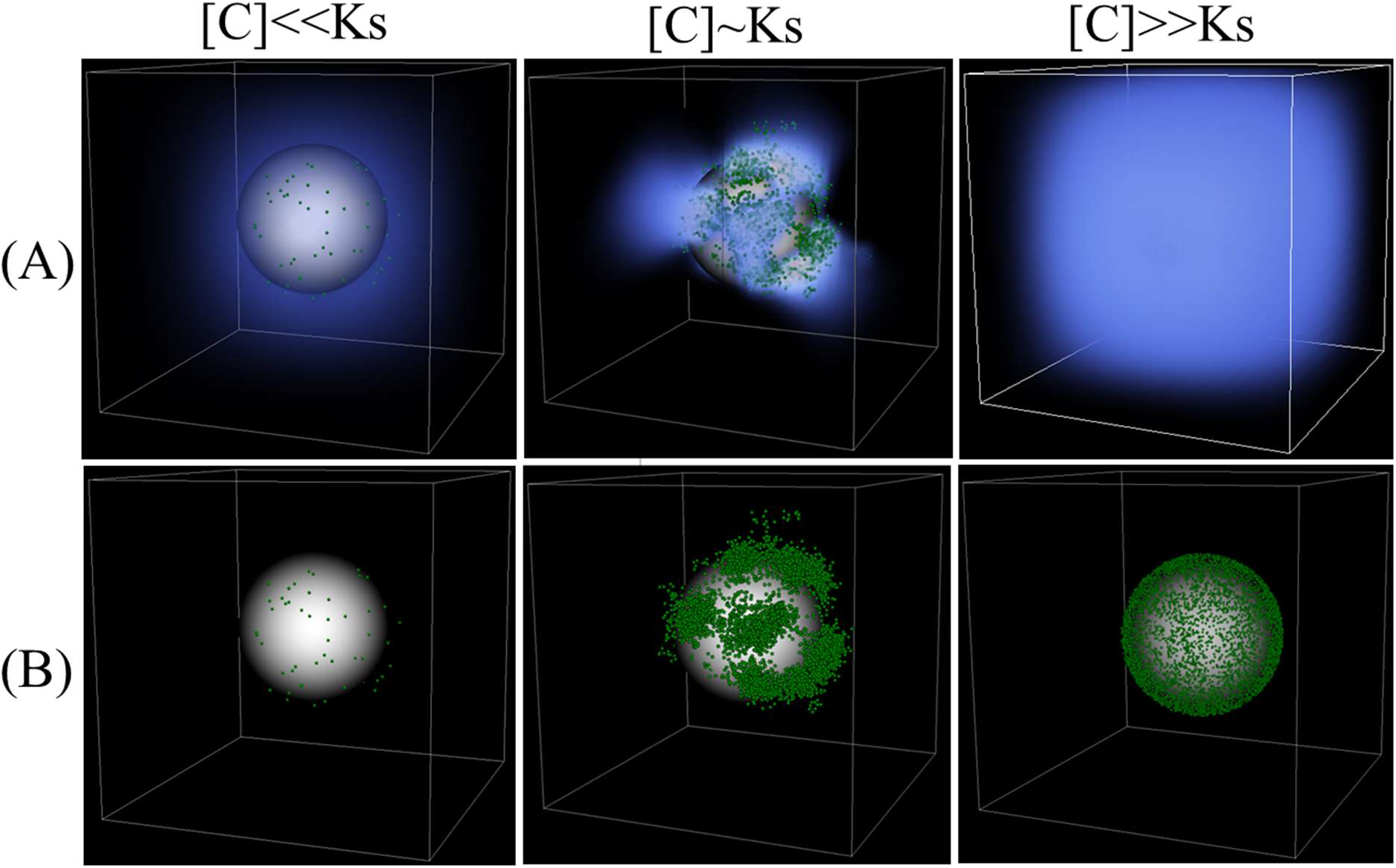
Spatial self-organization and cooperative growth allow degradation of recalcitrant organic particles. (A) Dissolved carbon concentration profile (red high and blue low concentrations) projected on microbial population assembly on the particle for three levels of particle lability (recalcitrant I: K_p_:1hr^−1^, semi-labile II: K_p_:100hr^−1^ and labile III: K_p_:1000hr^−1^). Green dots show individual cells and (B) only shows the bacterial cell colonization on the particle. For visualization purposes, carbon concentrations below a certain threshold are not shown (below 1% of maximum concentration). A particle size of 200 μm is considered and initial cell density was assumed to be 1000 cells.

**Figure S8.**
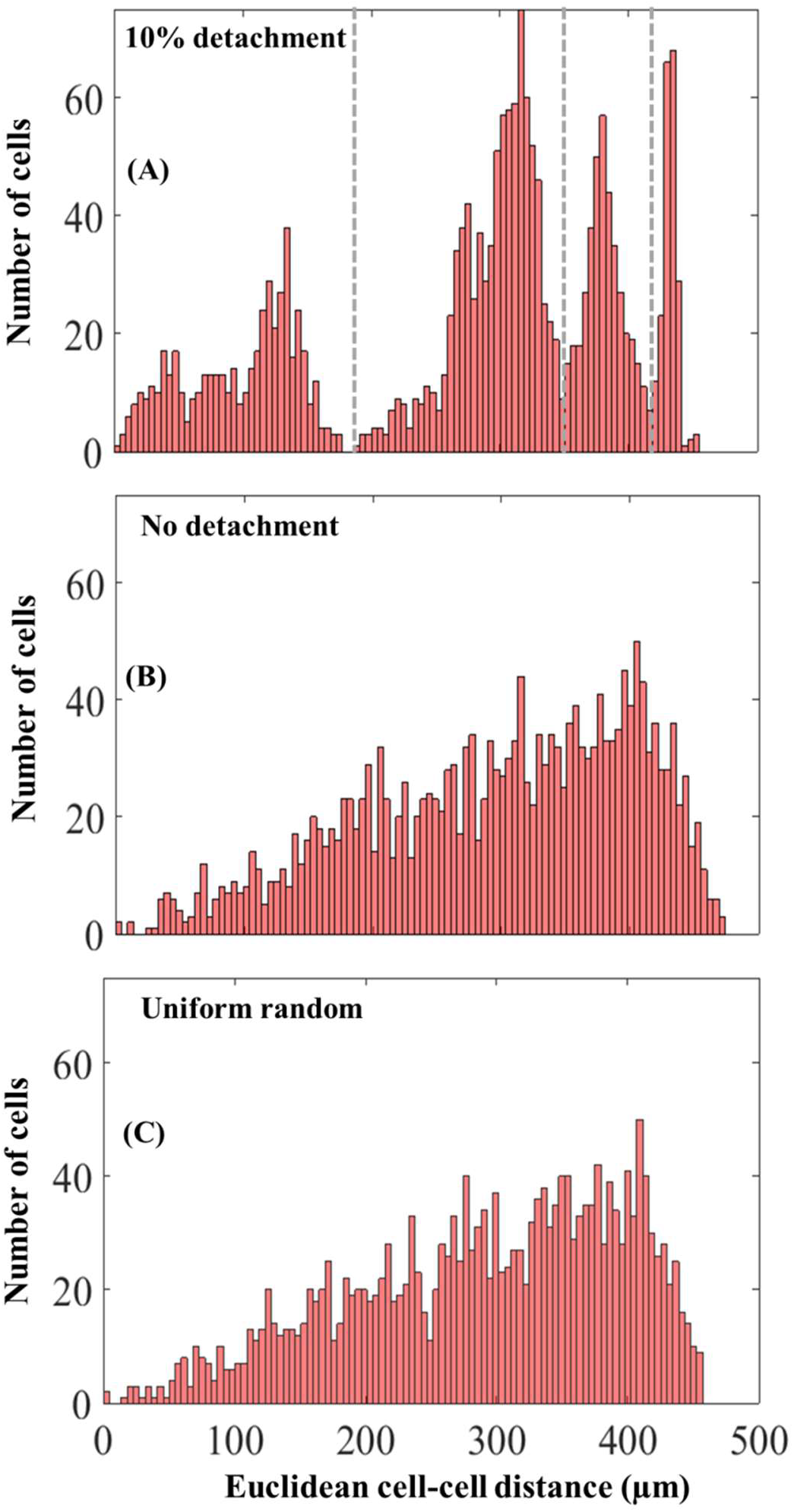
Detachment changes in the distribution of cell-cell distances within particle associated populations. Distribution shown for simulations with 10% detachment (A) or without detachment (B). The simulations were run for semi-recalcitrant particles (Kp=100 hr^−1^), corresponding to the simulations in Figure 2B with an initial cell density of 1000 and after 10 hours. The dashed grey lines separate single patches. (C) A hypothetical scenario with a uniform random distribution of bacterial cells on the particle surface is shown for comparison. In this comparison, the cell-cell distance was calculated between a reference single cell and randomly selected 2000 cells on the particle.

**Figure S9.**
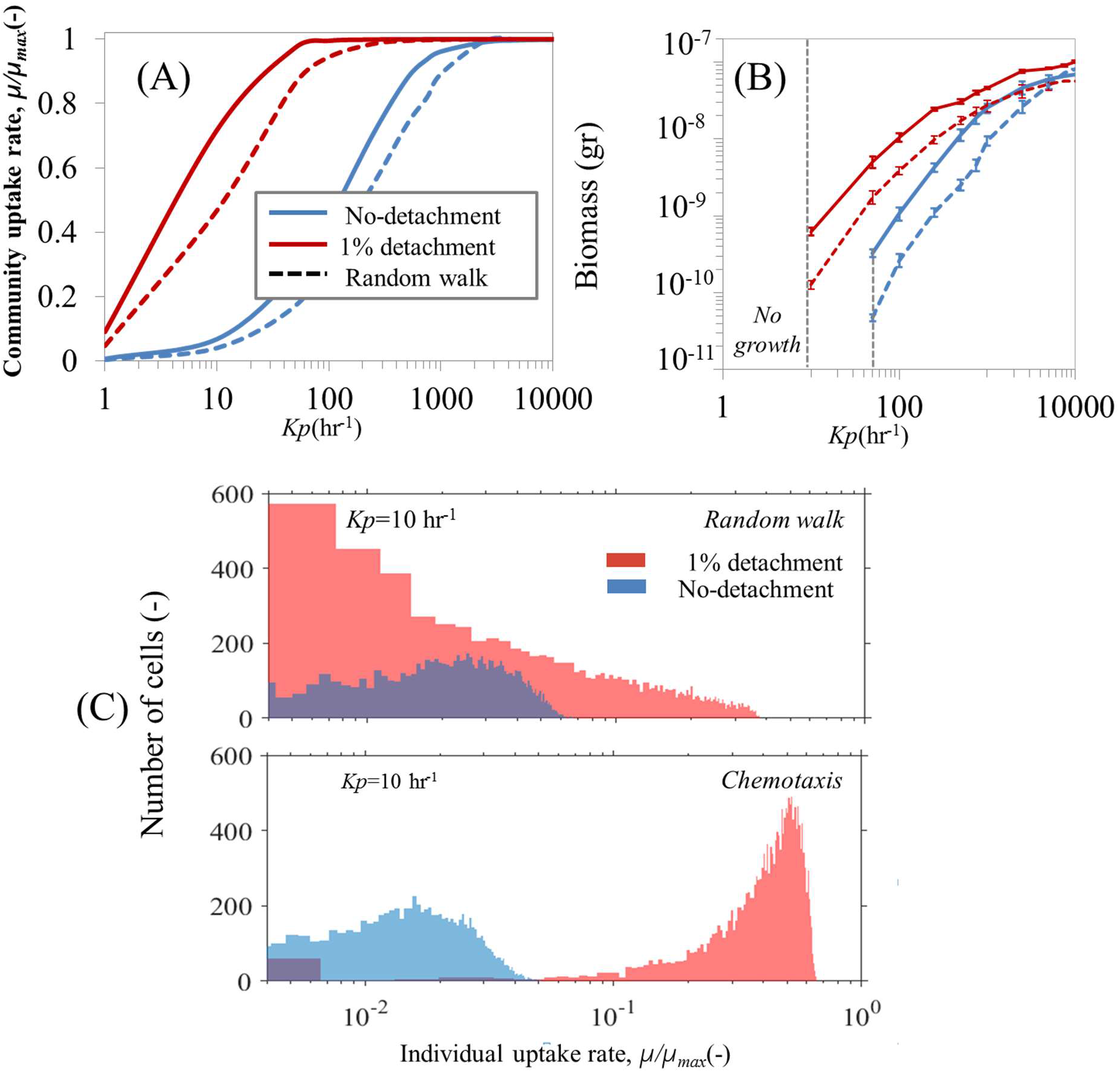
Microbial dispersal strategies regulate POM uptake efficiency from organic particle. (A) Effects of individual cells chemotactic behavior vs. random walk and attachment/detachment frequency to/from particles on Mean carbon uptake rates, represented as a function of particle lability (K_p_). The mean uptake rate is normalized by the maximum uptake rate of individual cells imposed by their physiological properties. (B) Biomass accumulation after 10hours. Grey dashed line indicates extinction zone of bacterial population (no biomass accumulation). (C) Histogram of individual cell uptake rates as represented by the number of cells colonizing the particle. The results are shown only for K_p_=10 hr^−1^. The initial cell number was 10000 cells and the particle size was set to 200 μm. The results are shown for simulations after 10 hours.

**Figure S10.**
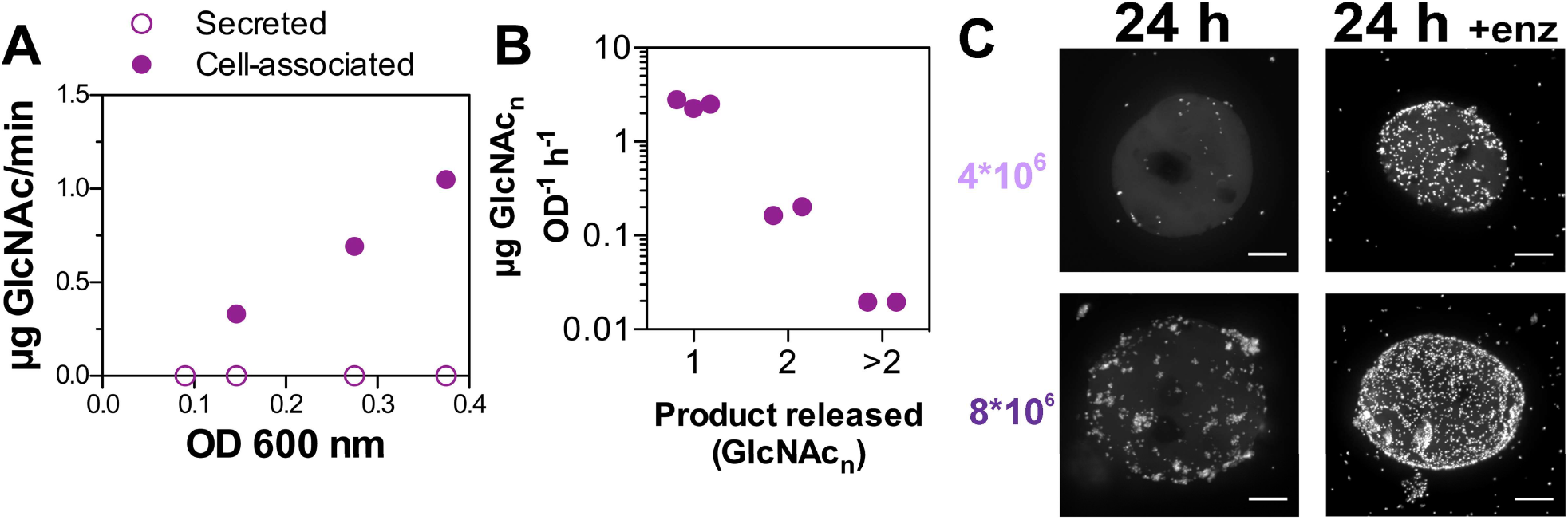
Chitinase production by psych6C06 is cell-associated, and exochitinase accounts for most of the activity. A) Activity of psych6C06 chitinases in culture supernatant, and cell-associated fractions. B) Activity of cell-associated chitinase for three different chitinase substrates (1, exochitinase-specific; 2, chitobiosidase-specific; 3, endochitinase specific). C) Representative images showing the effect of enzyme addition (Figure 3D) on the colonization pattern and density of psych6C06. Images were acquired 24 h after innoculation of chitin beads with 4*10^6^ cells/mL (light purple, below colonization threshold), or 8*10^6^ cells/mL (dark purple, above colonization threshold). Scale bars are 20 μm.

**Table S1.** Physiological parameters for microbial growth, metabolism and nutrient concentrations in the individual-based model.

| *Parameters* | *Values (Units)* |
| --- | --- |
| $\mu_{\max}$ : maximum growth rate ($hr^{-1}$) | 0.2-0.5* |
| $K_s$ : half saturation (mg/L) | 0.01-0.1* |
| $Y_{\max}$ : growth yield (gr dry mass/gr substrate) | 0.5* |
| cell maintenance | $0.1 \mu_{\max}^{\text{f}}$ |
| cell size ($\mu m$) | $1^{\text{f}}$ |
| $\rho$: cell density ($mg L^{-1}$) | $2.9 \times 10^5^{\text{f}}$ |
| $V_{ID}$ : cell velocity at bulk solution ($\mu m/s$) | $10^*$ |
| $V_u$ : median cell volume (fl) | $0.4^{\text{f}}$ |
| $K_{enz.}$ : enzyme production (gr enzyme/ gr biomass $hr^{-1}$) | $0.05^*$ |
| Tumbling frequency ($s^{-1}$) | $0.1^{\text{f}}$ |
| $X_0$: chemotactic sensitivity($mm^2/hr$) | $446^{\text{f}}$ |
| $^{\text{f}}$ (2) | |
| $^{\text{f}}$ (22) | |
| $^{\text{f}}$ (8) | |
| *Model assumption |  |

**Table S2.**
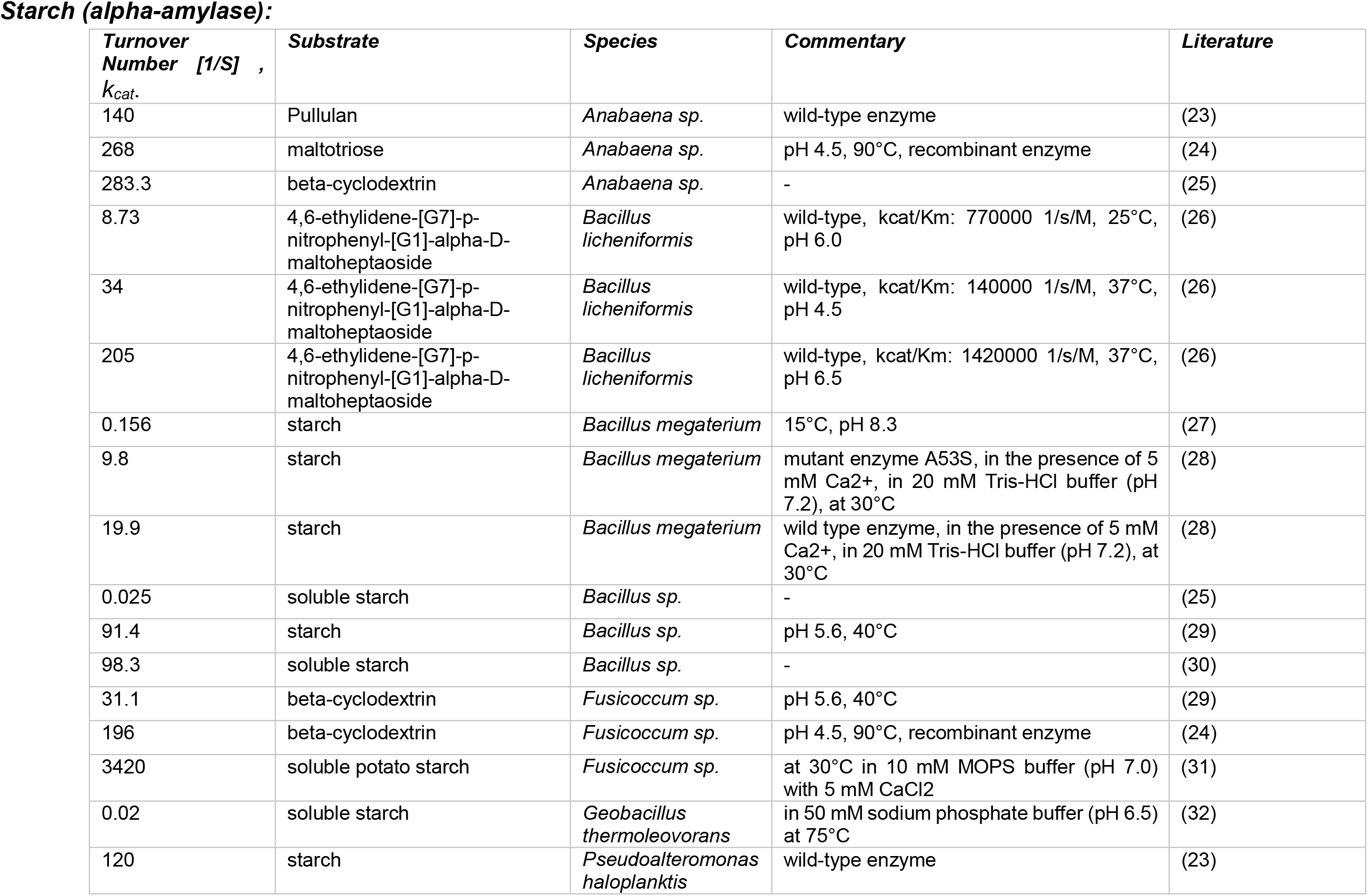
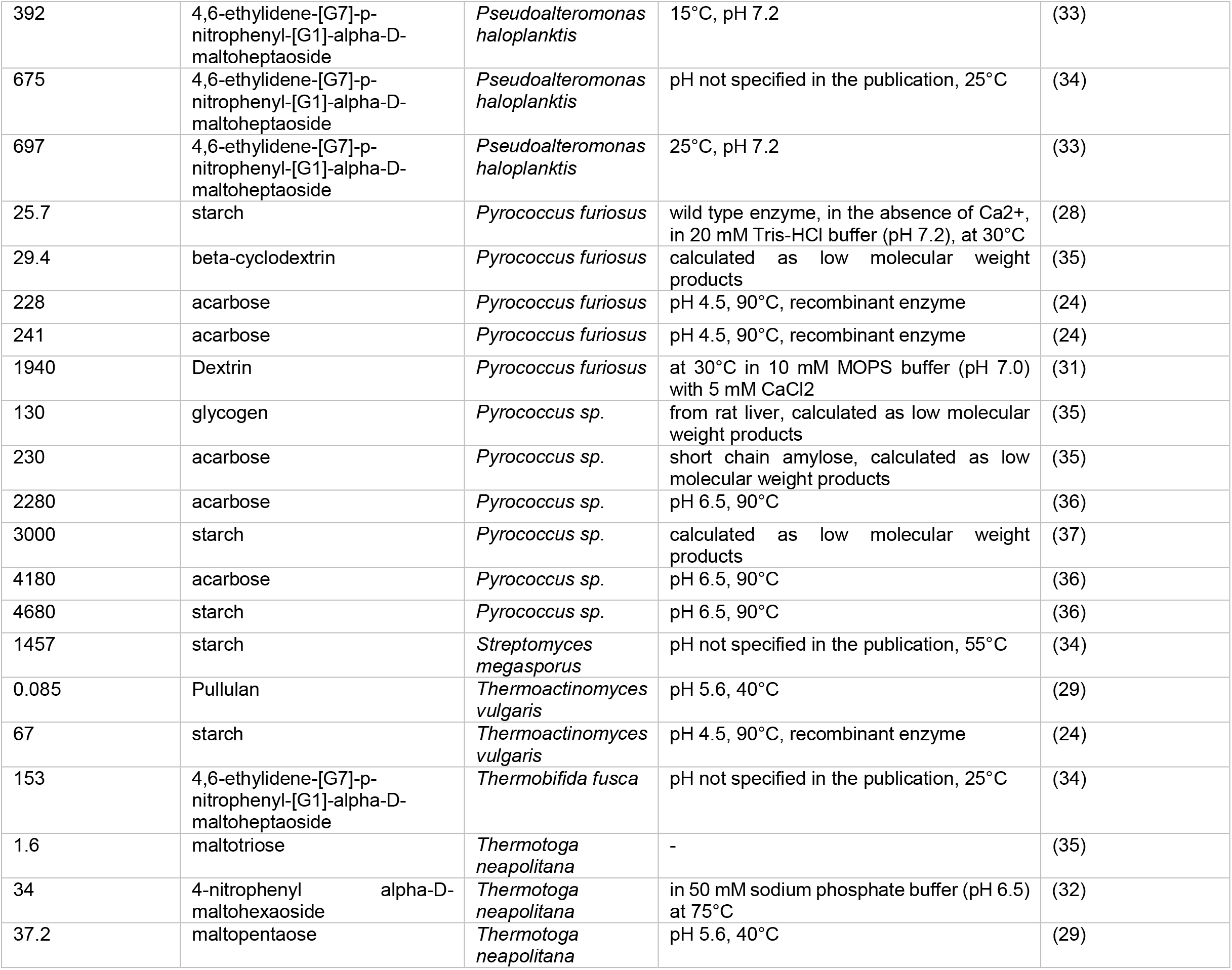
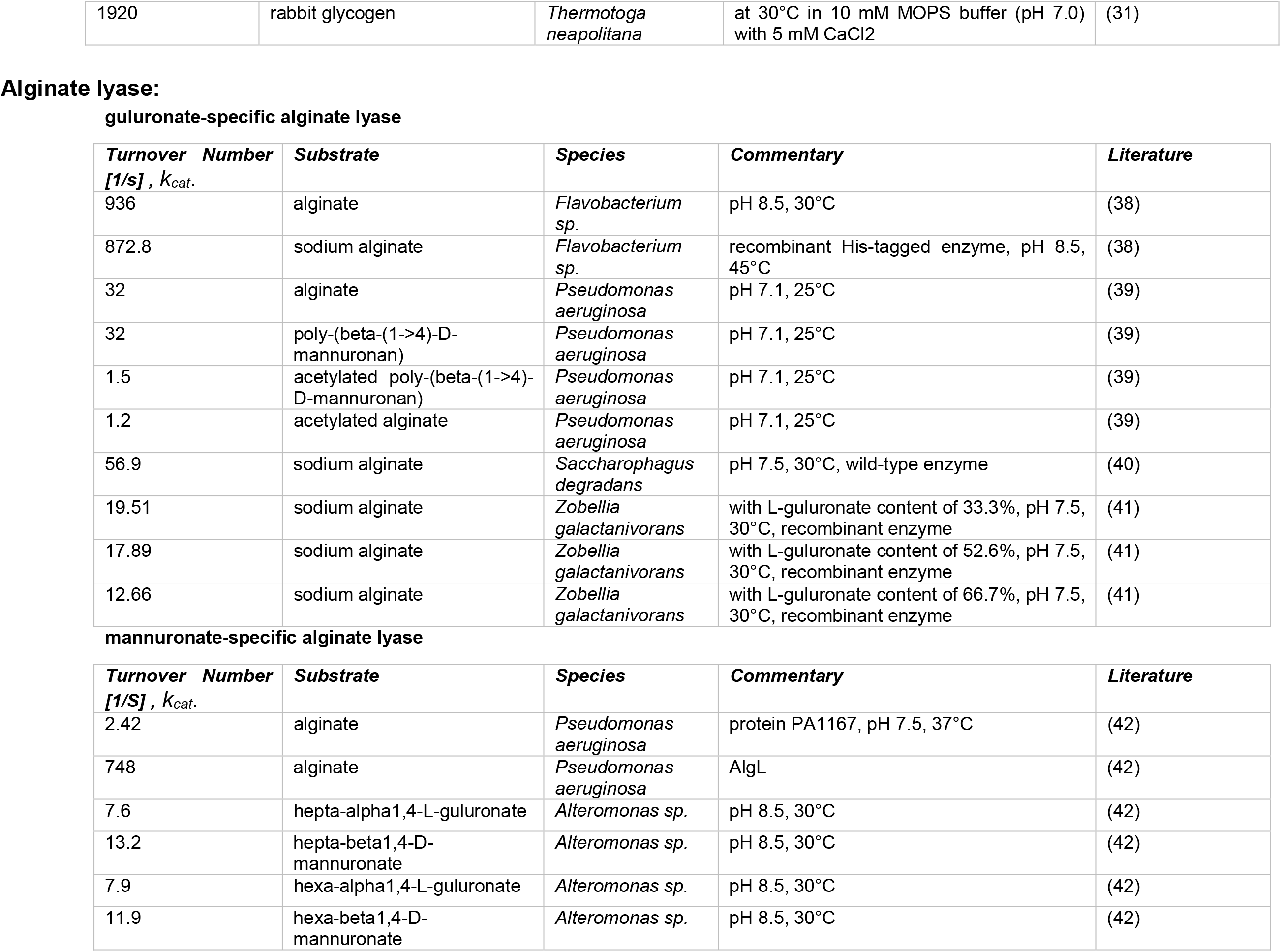
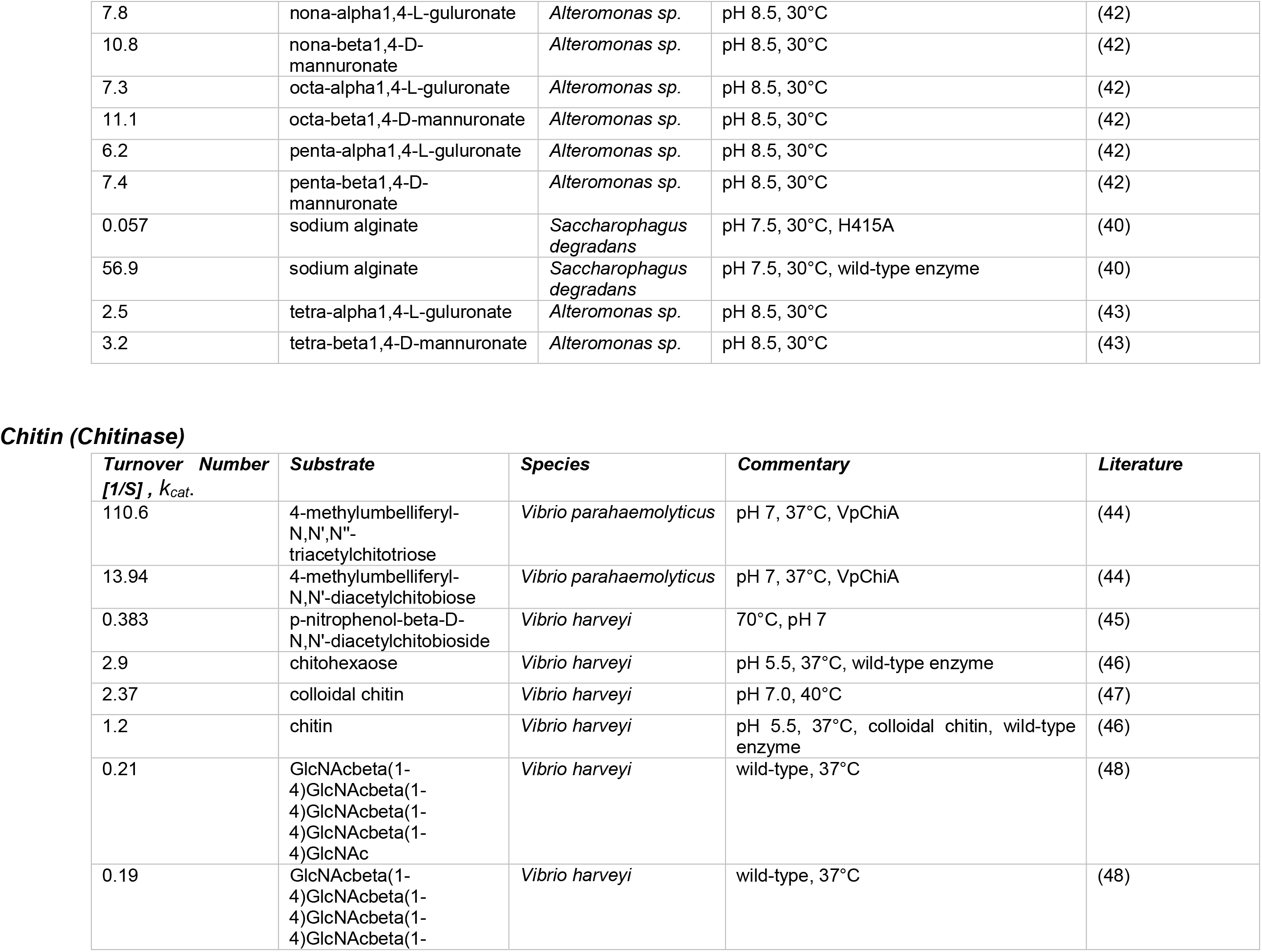
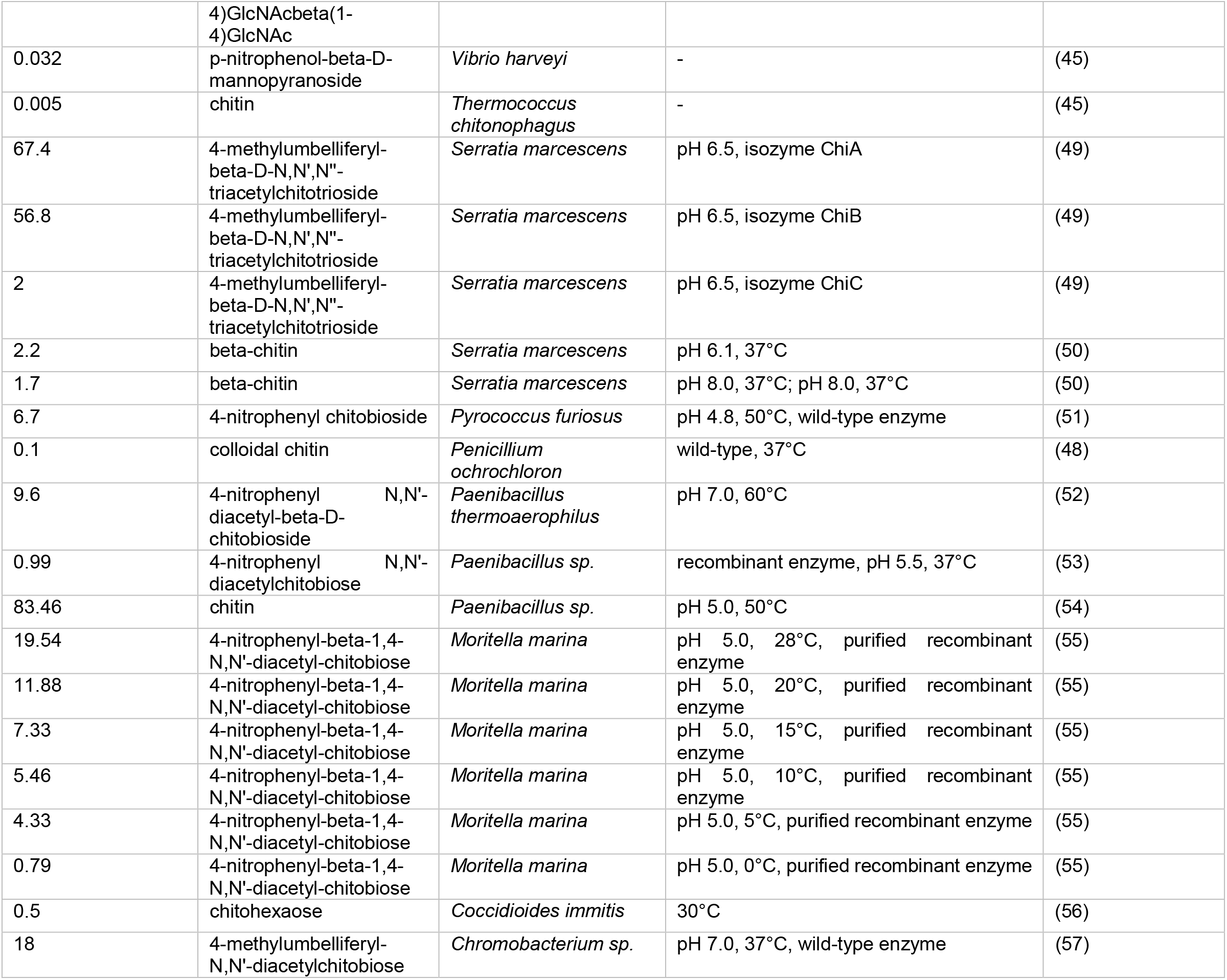
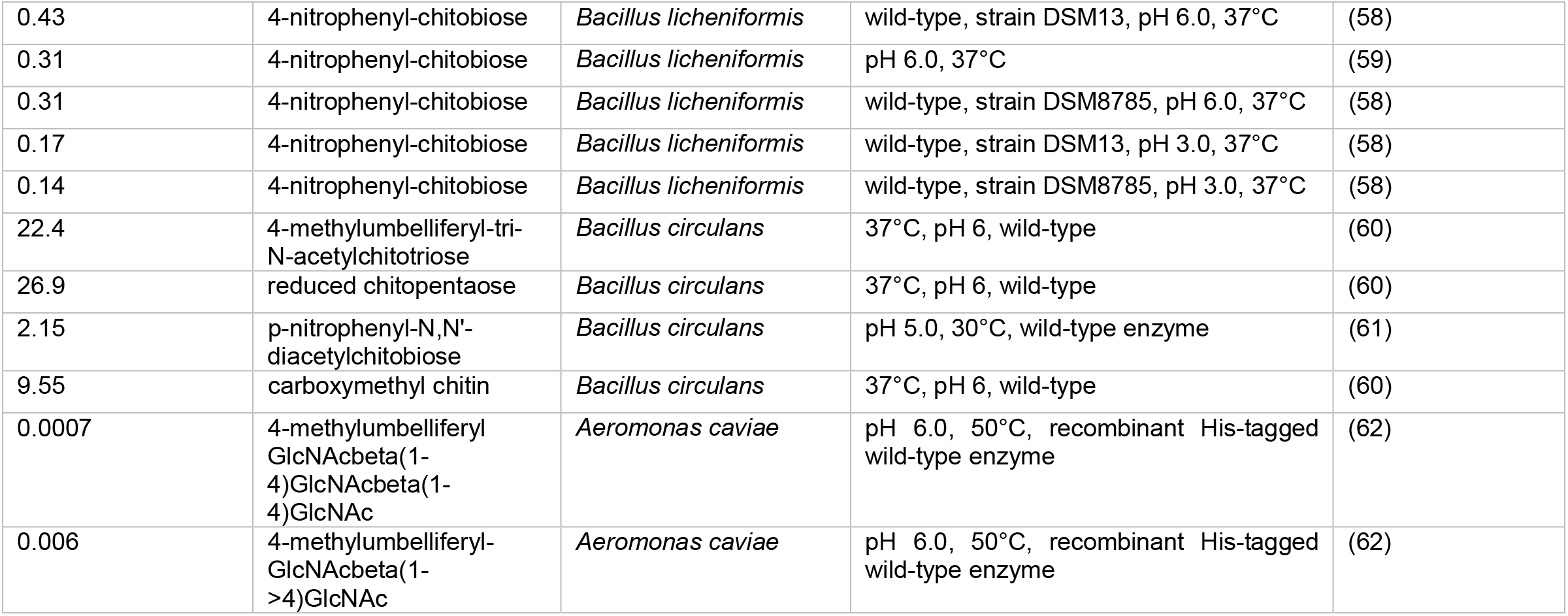
The distribution range of the catalytic activity of various enzymes kcat. as a measure for particle lability from natural polymeric carbohydrates (Chitin, Alginate, Starch). The data are for various bacterial species with their corresponding abiotic conditions (species name, substrates and environmental conditions)

**Table S3.**
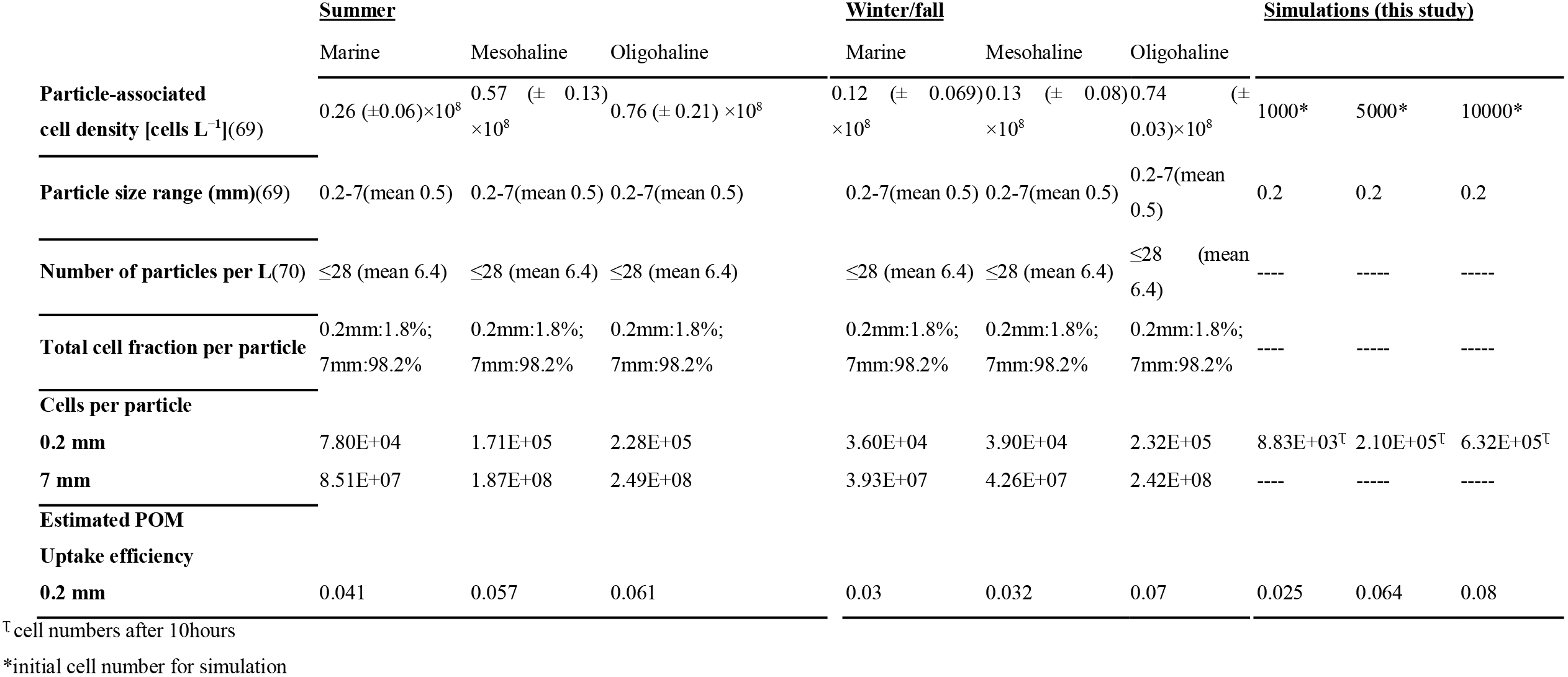
Particle associated cell density at different water solutions and seasons in the Baltic Sea. The data are extracted from Rieck et al., 2015 (69). The data on particle size range and number of particles per one litter of solution are used from field studies (70) to estimate the number of cells per single particles across observed size ranges. The simulations are performed for three initial cell densities for approximately the mean particle lability observed for natural biopolymers (K_p_~100hr^−1^, Figure 1B) and POM uptake efficiency is estimated after 10hours. The simulation data are used to predict uptake efficiency from natural marine snow.

